# Surface engineering enhances the therapeutic potential of extracellular vesicles following acute myocardial infarction

**DOI:** 10.1101/2022.01.31.478512

**Authors:** Kyle I. Mentkowski, Touba Tarvirdizadeh, Cody A. Manzanero, Lisa A. Eagler, Jennifer K. Lang

## Abstract

**Objectives:** The objective of the study was to assess the therapeutic efficacy of targeting remote zone cardiomyocytes with cardiosphere-derived cell (CDC) extracellular vesicles (EVs) in acute myocardial infarction.

**Background:** Cardiomyocyte (CM) cell death plays a significant role in left ventricular (LV) remodeling and cardiac dysfunction following myocardial infarction (MI). While EVs secreted by CDCs have shown efficacy in promoting cardiac repair in preclinical models of MI, their translational potential is limited by their biodistribution. We hypothesized that targeting therapeutic EVs to CMs would result in further reduction of cardiomyocyte (CM) cell death *in vivo* and improvement in cardiac function post-MI.

**Methods:** CDC-derived EVs were engineered to express a CM-specific binding peptide (CMP) on their surface and characterized for size, morphology, and protein expression. Mice with acute MI underwent delivery of human CDC EVs, CMP-EVs and placebo in a double-blind study. LVEF was assessed by echo at 1- and 28-days post-MI and tissue samples processed for assessment of EV biodistribution and histological endpoints.

**Results:** CMP-EVs demonstrated superior cardiac targeting and retention when compared with control EVs 24 hours post MI. While intramyocardial administered CDC-EVs improved LVEF compared with placebo at 4 weeks, mice treated with CMP-EVs demonstrated a significant improvement in LVEF compared with non-targeted EVs. Likely accounting for their augmented therapeutic efficacy, CMP-EVs demonstrated enhanced reduction of remote zone cardiomyocyte apoptosis.

**Conclusions:** Targeting CMP-EVs to CMs post-MI improved cardiac function compared with unmodified EVs demonstrating a strategy to further optimize therapeutic EV delivery to increase efficacy and decease off-target effects.

**Condensed Abstract:** Extracellular vesicles (EVs) offer several potential advantages over small molecule therapeutics for cardiovascular disease (CVD). However, their potential is limited by their biodistribution and lack of specificity. We engineered cardiosphere-derived cells (CDCs) to express Lamp2b fused to a cardiomyocyte specific peptide (CMP), generating EVs with increased cardiomyocyte uptake and cardiac retention. CMP-EVs enhanced cardiac function following acute MI and further reduced remote zone apoptosis when compared with unmodified CDC-EVs. This work highlights a role for targeting therapeutic EVs to cardiomyocytes following injury and serves as a proof-of-concept study for the utility of EV surface engineering in the treatment of CVD.

## Introduction

Cardiomyocyte cell death is a hallmark of cardiovascular disease (CVD) and contributes to the pathogenesis and progression of both ischemic and non-ischemic cardiomyopathy ^1^. While myocardial infarction and ischemia/reperfusion lead to acute, localized myocyte loss, global changes in cellular dynamics following cardiac injury can also potentiate chronic, diffuse myocyte cell death. Regardless of the inciting stimulus, progressive myocyte death contributes to a loss of functional myocardium. Given the lack of robust endogenous cardiomyocyte proliferation or cardiac regeneration in the adult human heart, development of targeted therapeutics to salvage at risk myocytes has the potential to alter the trajectory of heart disease.

Extracellular vesicles (EVs) have emerged as promising therapeutic vectors for cardiovascular disease owing to their natural properties of low immunogenicity, biocompatibility, and preservation of the therapeutic activity of their bioactive cargo ^2^. Recently, attention has focused on engineering the surface and cargo of therapeutic EVs to improve their biodistribution, pharmacokinetic properties, and therapeutic efficacy ^2,3^. EVs derived from cardiosphere-derived cells (CDCs) have demonstrated therapeutic efficacy in pre-clinical models of heart disease ^4–9^. Using a lentiviral knockdown strategy against neutral sphingomyelinase 2 (nSMase2), a crucial gene in ESCRT-independent EV secretion, we also defined a role of physiologically secreted human CDC-EVs on cardiomyocyte apoptosis ^10^. Prior studies have demonstrated intramyocardial delivery of CDC-EVs is superior to that of intracoronary delivery in reducing infarct size post-MI ^6^. We hypothesized that efficacy associated with intramyocardial delivery was related to higher myocardial retention and engineering the surface of EVs to target cardiomyocytes would further enhance their therapeutic efficacy.

We previously engineered CDCs to express Lamp2b, an EV membrane protein, fused to a cardiomyocyte specific peptide (CMP) ^11^. Extracellular vesicles isolated from engineered CDCs expressed CMP on their surface (CMP-EVs) and retained their native EV properties ^11^. Targeted EVs resulted in increased *in vitro* uptake by cardiomyocytes, decreased cardiomyocyte apoptosis in *in vitro* assays, and higher cardiac retention following intramyocardial injection when compared with non-targeted EVs ^11^. In this article, we sought to investigate if enhanced cardiac retention of therapeutic CDC-EVs translated to a functional benefit post-MI. We demonstrate that cardiomyocyte targeting of CDC-EVs enhances their ability to augment cardiac function. Furthermore, we provide evidence that targeting cardiomyocytes, not just the ischemic myocardium, has the potential to enhance systolic function post-MI via rescue of remote zone cardiac myocyte apoptosis. This work highlights a role for targeting therapeutic EVs to cardiomyocytes following injury and serves as a proof-of-concept study for the utility of EV surface engineering in the treatment of cardiovascular disease.

## Methods

Animal experiments were performed according to protocols approved by the University at Buffalo Institutional Animal Care and Use Committee. Human cardiosphere-derived cells were obtained from patients who consented to tissue use under protocols approved by the University at Buffalo Research Subjects Institutional Review Board. Methods were conducted in accordance with the relevant guidelines and regulations.

### Cell Culture

#### Cardiosphere-derived cells

Human cardiosphere-derived cells were generated as previously described under approved Institutional Review Board protocols ^10^. Briefly, endomyocardial biopsies were minced into small fragments, washed, and digested with type IV collagenase for 60 minutes at 37 °C. Explants were cultured on 20 μg/mL fibronectin-coated dishes. A layer of stromal-like cells and population of small, round, phase-bright cells migrated out to surround the explants. Once confluent, the cells surrounding the explants were harvested by gentle enzymatic digestion. These cardiosphere-forming cells were seeded at ∼1 × 10^5^ cells/mL on low attachment dishes in cardiosphere medium (20% heat-inactivated fetal calf serum, pen/strep 100 μg/mL, 2 mmol/L L-glutamine, and 0.1 mmol/L 2-mercaptoethanol in Iscove’s modified Dulbecco medium). Following 4–6 days in suspension culture, cardiospheres were harvested and plated on fibronectin-coated flasks forming monolayers of cardiosphere-derived cells. CDCs were characterized by flow cytometry and immunohistochemistry as previously described^7^.

#### HEK 293T cells

Human embryonic kidney (HEK) 293T cells were cultured in DMEM (Gibco) supplemented with 10% FBS (Corning) and pen/strep 100 μg/mL. All cells were incubated at 37°C in 5% CO_2_.

### Lentivirus generation and transduction

Lamp2-CMP lentivirus was generated as previously described ^11^. Briefly, HEK 293T cells were grown to 70% confluence and triple transfected with the lentiviral backbone pLenti-CMP and packaging plasmids pLP/VSVG (Invitrogen) and psPAX2 (AddGene) using Fugene HD (Promega). Viral supernatant was collected at 48 and 72hr and yielded a titer of at least 10^6^ GFP-transducing units/mL.

### Extracellular vesicle isolation and characterization

Extracellular vesicles were isolated from normal and transduced CDCs using an ultrafiltration method as previously described ^7^. Briefly, 70-90% confluent CDCs were cultured for 7 days in serum-free media. Supernatants were collected, passed through a 0.2 µm mesh filter, and EVs concentrated with an Amicon Ultra-15 centrifugal filter with a 10-kDa molecular weight cutoff. EV protein content was quantified with a Pierce BCA Protein Assay (Thermo Scientific). EVs were additionally characterized by size using Nanoparticle tracking analysis (NTA), surface charge (SZ-100, Horiba Scientific), morphology by cryo-TEM (FEI Vitrobot, Mark IV), and expression of known exosomal surface protein markers and markers of cellular contamination (Exo-Check Exosome Antibody Array, SBI) as previously described in detail ^11^.

### *Ex vivo* EV biodistribution studies

#### EV labeling

EV pellets were suspended in 500 µL of 1xPBS, mixed with lipophilic membrane dye DiR (Thermo) or DiD (Thermo) at a 1:10 volume ratio (dye/PBS), and incubated for 10 minutes at 37°C. Following incubation, 100 µL of ExoQuick-TC (System Biosciences) was added, inverted to mix and placed on ice for 30 minutes. The solution was centrifuged at 14,000xg for 3 minutes and the supernatant containing free dye aspirated and discarded. EVs were resuspended in PBS at a concentration of 10 µg/µL for *in vivo* use. DiR-labelling enabled whole organ bio-fluorescence measurements, while DiD-labelling enabled flow cytometry analysis of EV distribution.

#### Whole organ biofluorescence

2- and 24-hours following EV administration via intravenous or intramyocardial routes (described below), mice were euthanized by inhaled isoflurane overdose and subsequent cervical dislocation, and organs harvested and placed on ice in PBS in the dark for subsequent assessment of biofluorescence (IVIS Spectrum, Roswell Park Comprehensive Cancer Center). Images were taken and analyzed with Living Image using the following specifications: Excitation −745 nm, Emission −820 nm, FOV −13.4 cm, Auto Shutter Time. To calculate EV biodistribution as a percentage of dose, the radiant efficiency of each organ was divided by the sum of the bio-fluorescence signals from all organs.

#### Organ Dissociation

To investigate EV biodistribution on a single cell level, mice were euthanized 2-hours following intravenous injection of DiD-labelled EVs. Mice were immediately perfused with 10mL of ice-cold PBS followed by 10mL of pre-warmed collagenase buffer (0.05% Collagenase IV in HBSS). The lungs, liver and spleen were removed for subsequent dissociation. *Lungs* were transferred to a 60 mm petri dish and minced into <1mm pieces. Minced tissue was incubated with collagenase buffer for 25 minutes at 37°C and pipetted vigorously. The dissociate was passed through a 100 µm cell strainer, washed with wash buffer (PBS + 2% heat inactivated FBS + 2 mM EDTA), and centrifuged at 500 x g for 5 minutes to pellet the single cell suspension. The *liver* was transferred to a 60 mm petri dish, finely minced (<1mm pieces), incubated with collagenase buffer for 15 minutes at 37°C, and pipetted to further dissociate the tissue. The dissociate was passed through a 100 µm cell strainer, centrifuged at 500 x g for 5 minutes and re-suspended in 1X RBC Lysis Buffer (150 mM NH4Cl, 10 mM NaHCO3, 1 mM EDTA). RBCs were lysed for 5 minutes at room temperature and the lysis reaction stopped with the addition of 4 volumes of PBS. Remaining cells were centrifuged at 500 x g for 5 minutes. The *spleen* was dissociated by grinding the organ between two frosted glass microscope slides. The dissociate was collected with HBSS, passed through a 100 µm cell strainer and centrifuged at 500 x g for 5 minutes. Cells were then re-suspended in 1X RBC Lysis Buffer and incubated for 5 minutes at room temperature. The reaction was stopped by adding 4 volumes of PBS and centrifuging at 500 x g for 5 minutes.

#### Flow cytometry

Following single-cell dissociation, cell pellets were resuspended in PBS and incubated with Fc receptor blocker (TruStain FcX anti-mouse CD16/32) for 20 minutes on ice. Cells were incubated with rat CD68 (FA-11, ThermoFisher) at a 1:100 dilution for 1 h on ice, followed by a wash with Sort Wash Buffer (PBS + 2% FBS), and subsequent incubation with donkey anti-rat Alexa Fluor 555-conjugated secondary antibody (ThermoFisher) at a 1:500 dilution for 30 min on ice. Cells were analyzed for co-expression of CD68 and DiD-EV signal by flow cytometry (BD LSRFortessa).

### Murine model of acute MI

#### Thoracotomy and LAD ligation

Anesthesia was induced in adult 8-12wk C57BL/6J mice with a mixture of ketamine (100 mg/mL) and xylazine (20 mg/mL). A midline ventral cervical skin incision was performed and the muscles overlying the larynx and trachea were bluntly dissected and retracted to allow visualization of the intubation device through the exposed trachea. To intubate the mouse, the tongue was slightly retracted, and a 20-gauge 1” smooth needle was inserted through the mouth and larynx and into the trachea with care taken not to puncture the trachea or other structures in the pharyngeal region. The needle was advanced 8 to 10 mm from the larynx and taped in place to prevent dislodgment. Sterile lubricating drops were placed in the eyes. Mice were ventilated with room air supplemented with oxygen (1 L/min) at a stroke rate of 130 strokes/min and a tidal volume of 10.3 μL/g using a Inspira ASV small animal ventilator (Harvard Apparatus). The left chest was shaved, and a 1.5 cm skin incision made along the mid-axillary line. The left pectoralis major muscle was bluntly dissociated exposing the ribs. The muscle layers were retracted, and a left thoracotomy performed between the third and fourth ribs to visualize the anterior surface of the heart and left lung. The LAD was visualized emerging from under the left auricle and ligated by passing an 8-0 nylon suture under the LAD with two throws to secure the ligation. Successful infarction was verified by visualization of blanching of the distal myocardium and dynamic ECG changes. Using 6–0 polypropylene suture, the intercostal space was closed in an x-mattress pattern, followed by closure of the muscle layer in an interrupted suture pattern, and skin layers in a running mattress pattern. Mice received post-operative analgesia with 3.25 mg/kg SQ extended-release buprenorphine (Ethiqa XR, Fidelis Phamaceuticals) and were recovered on a heating pad.

#### Intramyocardial injection

Immediately following LAD ligation, mice were randomly selected to receive intramyocardial injections of PBS, unmodified EVs, or CMP-EVs. Equal numbers of male and female mice were selected across treatment groups to control for sex as a variable. Mice received a total of 100µg of EVs (or matched volume vehicle control) administered intramyocardially in two peri-infarct sites (10µl/injection) using a Hamilton syringe with a 30.5-gauge sterile beveled needle. The tip of the needle was bent at a 45° angle and introduced into the left ventricular myocardium. The solution in the syringe was slowly injected and local tissue blanching verified, indicating intramyocardial delivery vs. LV cavity administration. The syringe was held in place for an additional 3–5 seconds then withdrawn.

#### Intravenous injection

Following extubation, mice received an IV tail vein injection of 100µg of EVs in 100µL of PBS (or matched volume vehicle control).

### Echocardiography

Transthoracic echocardiography (Vivid E9 with GE Ultrasound i13L Intraoperative Epicardial Probe) was performed on mice 2 days and 28 days post-MI. Depilatory cream was applied to the chest to remove hair to reduce imaging artifacts. Mice were placed on a heated surgical platform (Rodent Surgical Monitor+, Indus Instruments) to maintain appropriate body temperature and noninvasively monitor heart rate, ECG, respiration rate, and SpO_2_ during echocardiography. Left ventricular ejection fraction was calculated from M-mode images obtained in the parasternal short axis mid papillary view. Heart rate was maintained between 500 and 550 bpm during image acquisition.

### Tissue Processing

Animals were euthanized at the terminal endpoints of the study and hearts perfused with KCL followed by PBS. For subsequent immunohistochemistry, a portion of the hearts were fixed for 30 min in 4% paraformaldehyde (PFA), cryopreserved in sucrose gradient (6, 15%) for 24 hours, snap frozen in OCT (optimal cutting temperature) compound (Tissue-Tek) and sectioned transversely at 12µm thickness using a Leica cryostat. For subsequent trichrome staining, the remaining portion of the hearts were fixed in 10% normal buffer formalin (NBF) overnight at room temperature, sectioned transversely into 1mm thick slices, and paraffin embedded apex to base. Paraffin blocks were further transversely sectioned at 5µm thickness and stained with Trichrome by the University at Buffalo Histology Core.

### Immunohistochemistry

In general, OCT sections were permeabilized for 30 minutes with 1% Triton X (ThermoFisher) and blocked for 1 hr with a solution containing 0.5% TritonX-100 and 5% donkey serum (ThermoFisher). Primary antibodies used were rat CD68 (FA-11, 1:100; ThermoFisher), rabbit cardiac troponin I (1:50; Proteintech), F-actin (1:200; Rhodamine Phalloidin, ThermoFisher) and rabbit cleaved caspase-3 (1:200; Asp175, Cell Signaling). Alexa Fluor 488-, 555-, and 647-conjugated secondary antibodies (ThermoFisher) were used at 1:500. Tissue sections were counterstained with DAPI (ThermoFisher). Images were captured at 20x magnification using an EVOS FL Auto Imaging System and an Olympus IX83 microscope with wide field fluorescence and at 63x magnification using a Leica TCS SP8 Confocal microscope. Macrophage density was calculated using a Fiji thresholding technique as the CD68+ area/total DAPI+ area. The percentage of apoptotic cells was calculated using Fiji as cleaved caspase-3^+^ cells/total cells. Corrected total fluorescence was calculated using Fiji as: integrated density – (total area*background integrated density) and displayed as a ratio of CDC-EV vs. CMP-EV total fluorescence. Percent DiD^+^ cardiac area was calculated as: DiD^+^ area/total cardiac area. Paraffin embedded trichrome stained sections were used for quantification of interstitial fibrosis and cardiomyocyte size and nuclear density. Cardiomyocyte area was calculated on cardiomyocytes with a short-axis orientation at 40x magnification. The criteria for short axis were a circular shape with a discernable nucleus. Cardiomyocyte nuclear density was calculated as the number of myocyte nuclei per mm^2^ using multiple compact fields with short-axis myocytes at 40x magnification. Interstitial fibrosis was determined using a Fiji thresholding method to compare the amount of blue to red staining. All quantification was performed by an investigator blinded to the identity of the samples.

### Statistical Analyses

Data were analyzed using GraphPad Prism software and are represented as mean ± SEM. Experiments comparing differences between two groups were analyzed by a two-tailed student’s t-test. Any comparison of more than two groups with one variable was analyzed by One-Way ANOVA with Tukey’s post hoc test. The cutoff for statistical significance was established as p<0.05.

### Data Availability

All data associated with this study are included in the published manuscript.

## Results

### CMP surface engineering alters EV cardiac biodistribution and pharmacokinetics following MI

To determine if cardiac targeting and retention of extracellular vesicles could be improved post-MI through surface expression of a cardiomyocyte-specific peptide, we studied the *in vivo* biodistribution and pharmacokinetics of previously characterized CMP-EVs ^11^. After acute MI, we performed blinded intramyocardial or intravenous injections of DiR-labeled EVs followed by *ex vivo* whole-organ fluorescence imaging at 2 and 24 hours (n=5-9 mice per group; CMP-targeted exosomes, unmodified EVs, PBS) (**Fig. 1a**). Following intravenous administration, CMP-EVs exhibited enhanced targeting to the heart post-MI when compared with unmodified EVs at both 2 hours (0.24 ± 0.6 vs. 0.07 ± 0.02) and 24 hours (0.44 ± 0.07 vs. 0.26 ± 0.04) (mean biodistribution as a % of dose ± SEM, *p<0.05 using paired two-tailed t-test) (**Fig. 1b,c**). This effect was specific to the post-cardiac injury milieu, as the ability of systemically administered CMP-EVs to target the heart was abrogated when they were injected into healthy mice, with both DiR-labeled CMP and control EVs demonstrating no detectable level of cardiac fluorescence above that of PBS control at 2 hours (p>0.05 using paired two-tailed test) (**Supplementary Fig. 1**).

**Figure 1.**
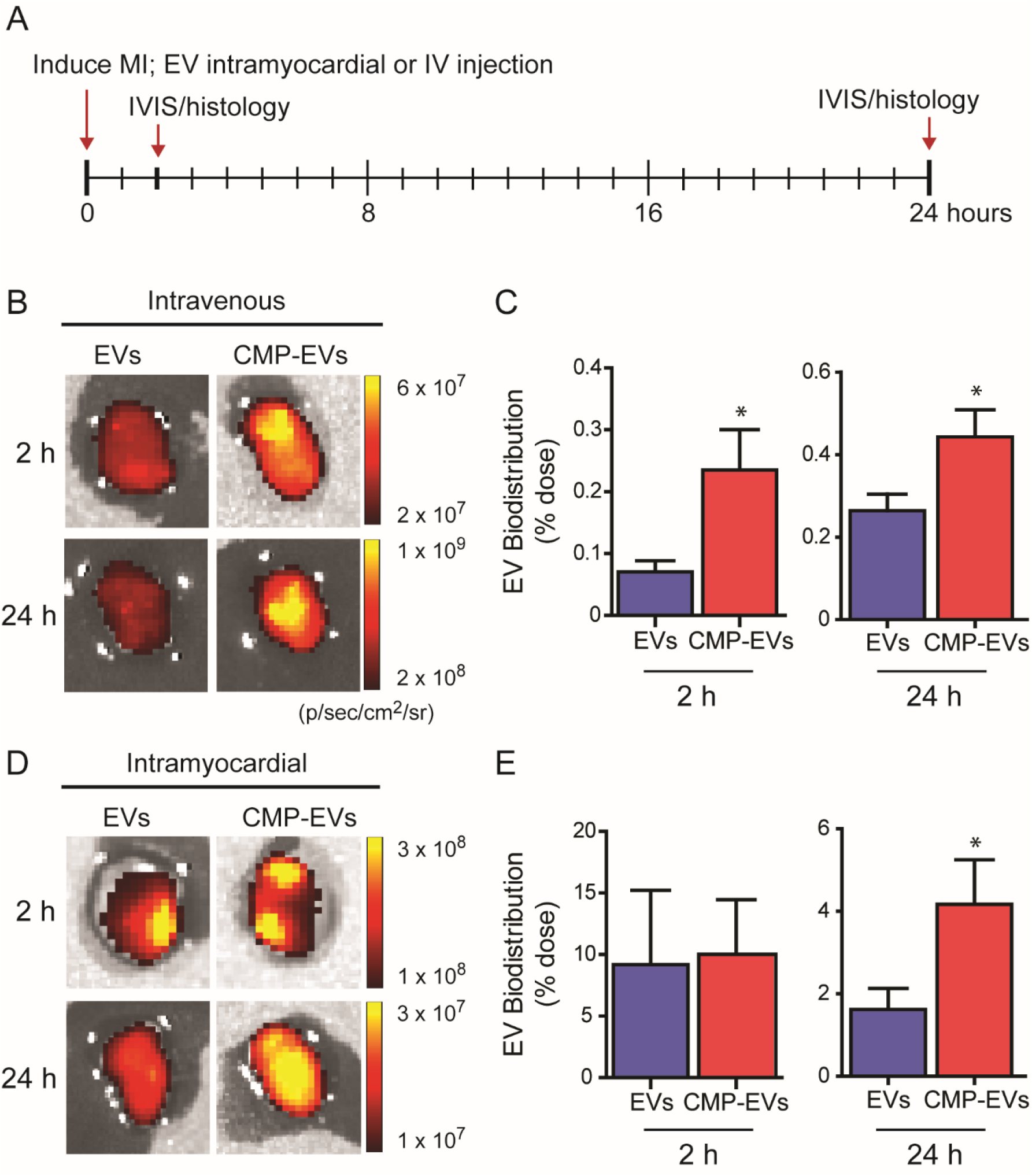
EV cardiac biodistribution and pharmacokinetics post-MI. **(A)** Schematic depicting the timeline of MI induction, EV administration and biodistribution/PK analysis endpoints. **(B)** Representative IVIS images of the heart following MI and intravenous EV administration demonstrates enhanced cardiac targeting of CMP-EVs vs unmodified EVs at 2 and 24 hours. **(C)** Normalized DiD-stained EV biofluorescence signal as a percentage of total intravenous injected dose at 2 and 24 hours. **(D)** Representative IVIS images of the heart following MI and intramyocardial EV administration demonstrate enhanced cardiac retention of CMP-EVs vs unmodified EVs at 24 hours. **(E)** Normalized DiD-stained EV biofluorescence signal as a percentage of total intramyocardial injected dose at 2 and 24 hours. n = 5-9 mice per group. *p<0.05 using a two-tailed student’s t-test. All data are represented as mean ± SEM.

When EVs were administered by intramyocardial injection following MI, cardiac biodistribution as a percentage of dose increased by approximately 10-fold when compared with IV administration. While CMP-EVs and unmodified EVs exhibited matched levels of cardiac retention 2 hours after intramyocardial injection (p>0.05 using paired two-tailed t-test), CMP-targeted EVs demonstrated higher cardiac retention at 24 hours when compared with non-targeted EVs (4.17 ± 1.08 vs. 1.62 ± 0.51) (mean biodistribution as a % of dose ± SEM, *p<0.05 using paired two-tailed t-test) (**Fig. 1d,e**).

In target tissues, EV localization was highest in the liver, lungs and spleen regardless of administrative route or time point assayed (**Supplementary Fig. 2**). When directly comparing CDC EV and CMP-EV uptake in each organ, we saw no significant differences between EV groups in the lungs, liver, spleen, gastrointestinal tract (GI), kidneys, brain, or bone marrow (**Supplementary Figs. 3-9**) (n=5-9 mice per group, p>0.05 using one way ANOVA). To investigate the effect of CMP-EV surface engineering on macrophage EV clearance following systemic administration, DiD-stained EVs were injected via an IV route and the lungs, liver and spleen removed and dissociated to a single cell suspension at 2 hours. CD68^+^ macrophages in the liver, lungs and spleen demonstrated the highest percentage of EV internalization, with no significant difference in DiD^+^/CD68^+^ cells between CMP-targeted and control EVs (**Supplementary Fig. 10**) (n=3 mice per group, *p<0.05 using one-way ANOVA). In summary, while CMP surface engineering did not enhance or reduce EV uptake by the reticuloendothelial system, CMP-EVs demonstrated enhanced cardiac targeting following systemic intravenous injection and augmented cardiac retention following intramyocardial administration.

### CMP targeting confers a higher area and density of EV cardiac retention post-MI

To further examine the effects of surface engineering on CDC EV cardiac biodistribution, we analyzed left ventricular (LV) transverse sections from intramyocardially administered DiD-EV groups at 24 hours. While unmodified CDC EVs were retained solely along the track sites of intramyocardial injection (**Fig. 2a**), CMP-EVs were seen in pockets throughout the LV with pronounced localization across the entire anterior wall (**Fig. 2b**). Compared with unmodified EVs, CMP-targeted EVs distributed and were retained across a larger total area 24 hours following intramyocardial injection (10.30 ± 2.10 vs. 4.27 ± 0.35%, %DiD+ cardiac area) (**Fig. 2a-c**). Furthermore, within regions of DiD+ EV localization, CMP-EVs demonstrated a higher ratio of total fluorescence in comparison with unmodified EVs (3.3 ± 0.6 vs. 1.0 ± 0.3), corresponding to a higher CMP-EV density within areas of EV retention (**Fig. 2d**) (n = 3 mice per group, mean ± SEM, *p<0.05 using paired two-tailed t-test).

**Figure 2.**
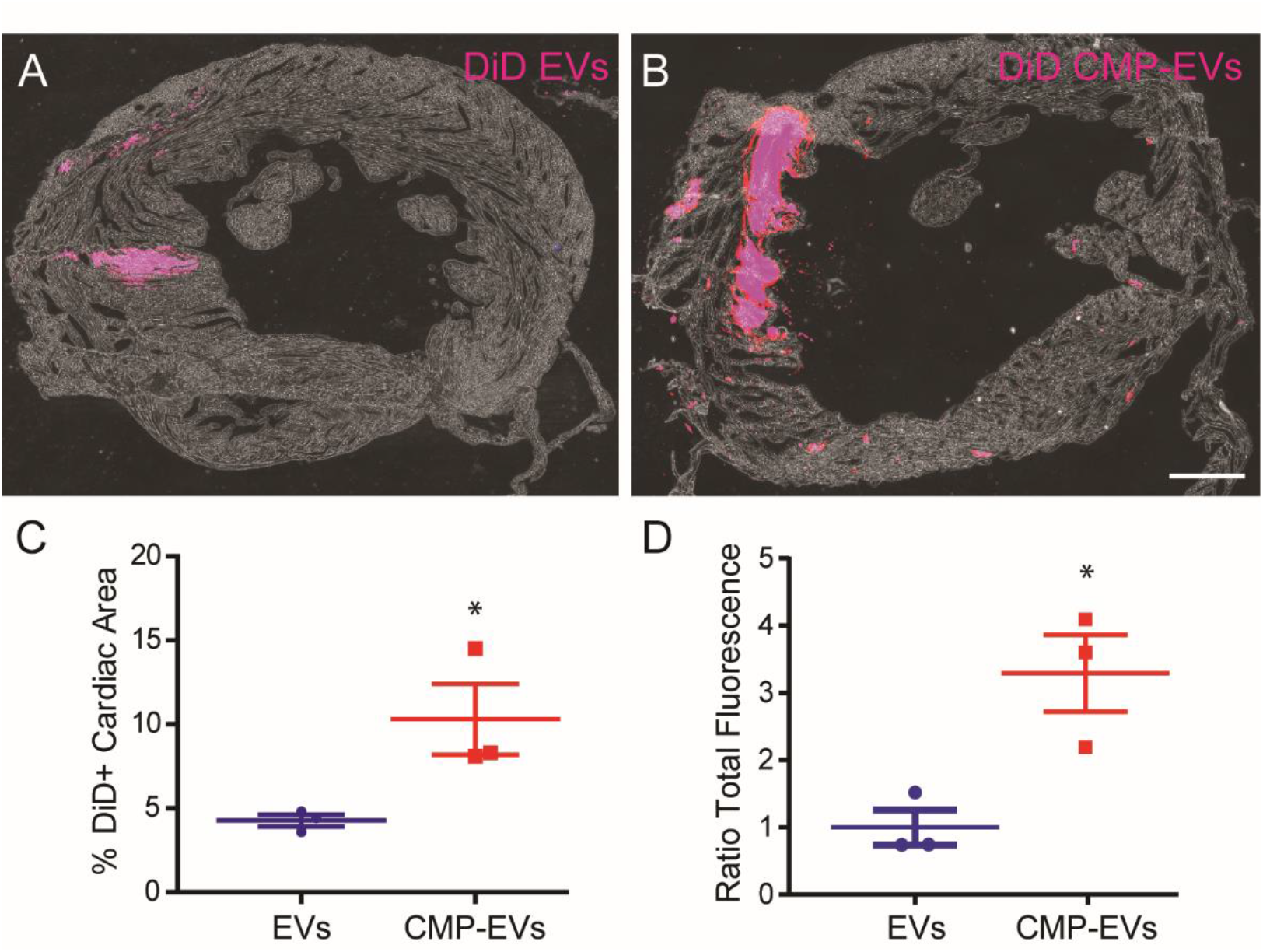
CMP-EVs demonstrate enhanced cardiac retention by area and density post-MI. Representative transverse left ventricular (LV) sections highlight differences in DiD-labeled **(A)** unmodified EV and **(B)** CMP-EV cardiac retention following intramyocardial injection 24 hours post-MI. **(C)** CMP-EVs are retained across a greater LV area (as quantified by % DiD^+^ area/total LV area) and **(D)** occupy a greater density within areas of EV retention (ratio of total fluorescence) when compared with unmodified EVs. n = 3 mice per group. Scale bar = 200 µm. *p<0.05 using a two-tailed student’s t-test. All data are represented as mean ± SEM.

### CMP surface engineering promotes EV localization and internalization by cardiomyocytes

To verify EV uptake into recipient cells, we used confocal microscopy to further examine the *in vivo* cellular distribution patterns of intramyocardially delivered DiD-labeled CDC EVs and CMP-EVs. Within regions of EV localization, CMP-EVs demonstrated enhanced cardiac retention and cardiomyocyte localization when compared with unmodified EVs (**Fig. 3a**). When confocal images were captured with matched exposure times and illumination intensity, Z-stack confocal analysis demonstrated enhanced CMP-EV internalization by TnI^+^ cardiomyocytes when compared with CDC EVs (**Fig. 3b**). Together, these data highlight the regional and cell-specific effects of CMP-targeting on EV cardiac biodistribution, retention and cellular uptake.

**Figure 3.**
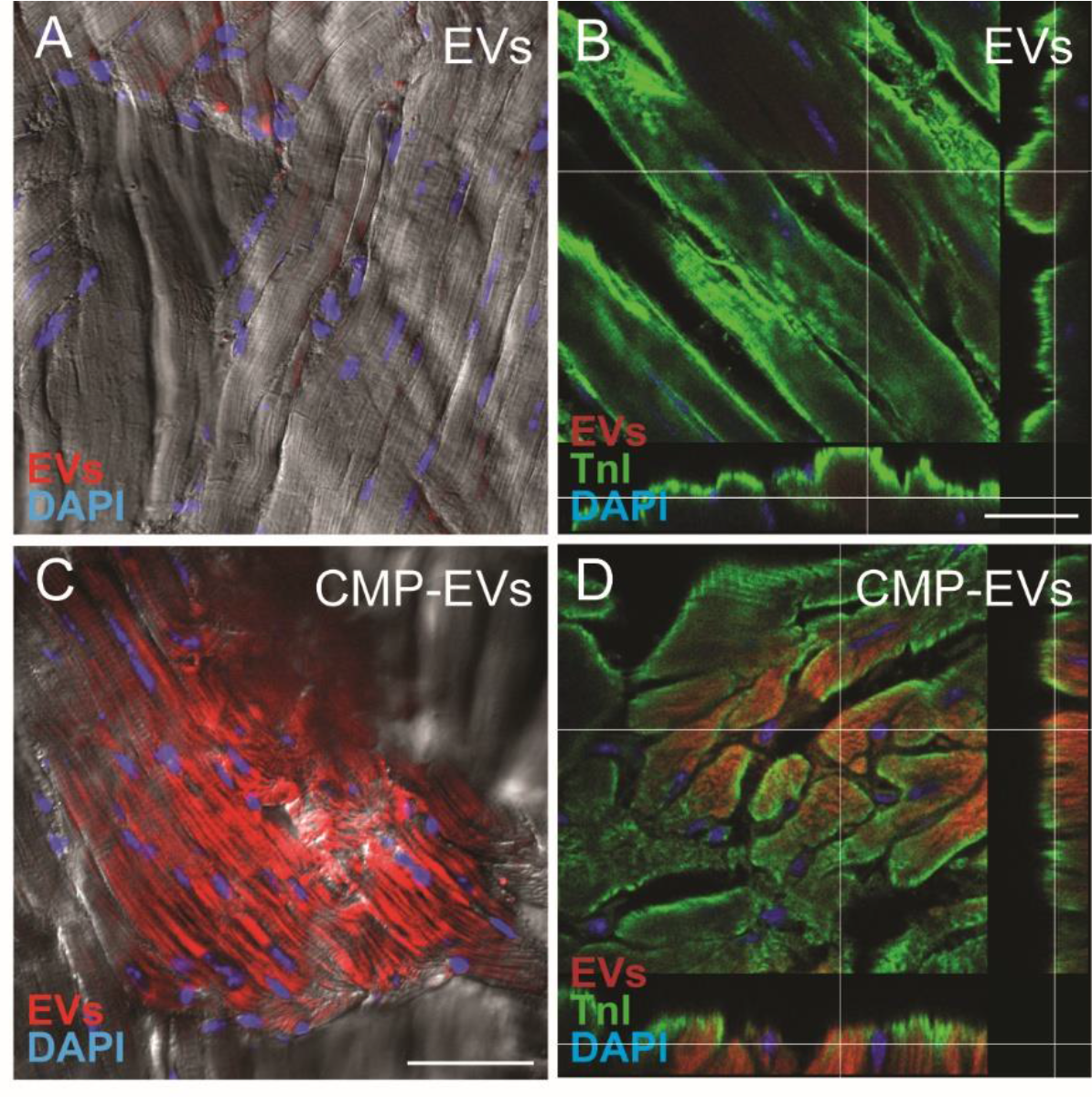
Confocal analysis of cardiomyocyte EV internalization. **(A,C)** Cardiomyocyte localization and **(B,D)** cellular uptake of DiD-labeled EVs delivered through intramyocardial injection were observed using confocal microscopy. CMP-EVs demonstrated **(C)** enhanced cardiac retention and **(D)** unaltered internalization by cardiomyocytes when compared with **(A,B)** non-targeted EVs. Scale bar = 20 µm.

### Cardiomyocyte targeting of CDC EVs enhances their potential to augment cardiac function and reduce fibrosis

To assess the therapeutic efficacy of CMP-EV targeting in post-infarct myocardium, CDC EVs and CMP-EVs were administered by intramyocardial injection at the time of MI and followed with serial echocardiography to assess changes in left ventricular ejection fraction (**Fig. 4a**). At 2 days post-MI, both EV groups and PBS vehicle control demonstrated equivocal left ventricular ejection fraction (LVEF; PBS: 45% ± 1%, EVs: 43% ± 2%, CMP-EVs: 41% ± 1%) (**Fig. 4b**). At 28 days post-MI, LVEF was significantly increased by CDC EV therapy vs PBS control (55% ± 2% vs. 44% ± 2%) and further augmented by CMP-EVs (64% ± 5%) (n=6-8 mice per group, mean± SEM, *p<0.05 using one-way ANOVA followed by Tukey’s post-hoc test) (**Fig. 4b**). Intramyocardial administration of CMP-EVs lead to a significant decrease in infarct size compared with PBS-treated controls (12% ± 5% vs. 28% ± 4%) (**Fig. 4c**) when analyzed by endocardial length-based measurement of Masson’s Trichrome stained mid LV cavity sections from PBS control, CDC EV and CMP-EV treated hearts at 28 days post-MI (n= 3-5 mice per group, mean ± SEM, *p<0.05 using one-way ANOVA followed by Tukey’s post-hoc test) (**Fig. 4d-f**). In summary, enhanced cardiac retention and cardiomyocyte uptake of CMP-EVs translated to improved systolic function and decreased infarct size following AMI.

**Figure 4.**
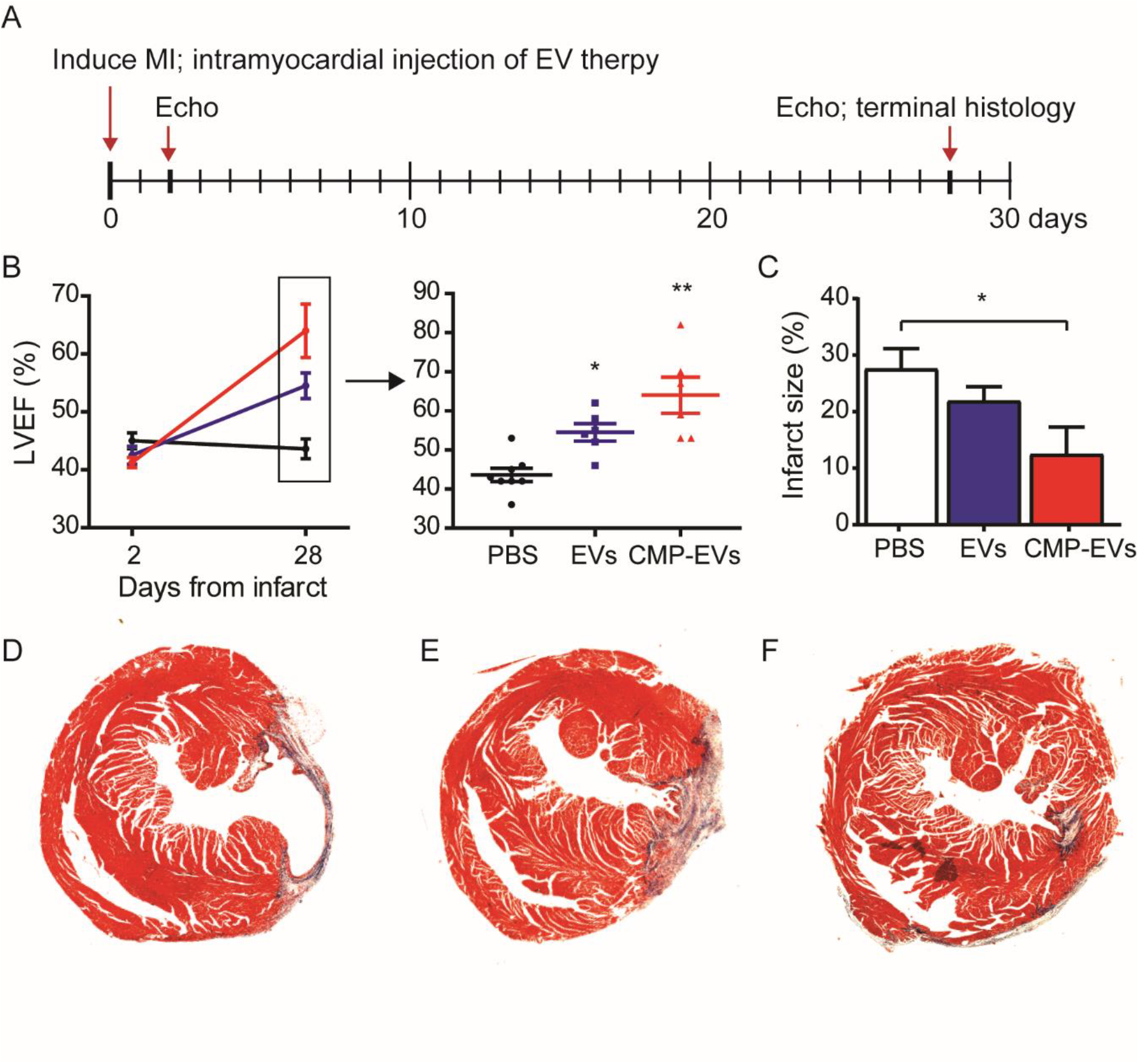
Cardiac function following EV treatment post-MI. **(A)** Representation of experimental design showing EV administration and imaging endpoints in mice after myocardial infarction. **(B)** The increase in left ventricular systolic function in CDC EV hearts compared with PBS control is further augmented by CMP-EVs at 28-days post-MI. n=6-8 mice per group. *p<0.05, **p<0.01 using a one-way ANOVA with Tukey’s post-hoc test. **(C)** Infarct size in CMP-EVs compared to CDC-EVs and PBS treatment groups 28 days after injury. n=3-5 mice per group. * p<0.05 using a one-way ANOVA with Tukey’s post-hoc test. Representative LV mid-cavity trichrome stained sections of **(D)** PBS, **(E)** CDC-EV, or **(F)** CMP-EV treated hearts at the terminal study endpoint. All data are represented as mean ± SEM.

### Remote zone interstitial fibrosis is significantly reduced by CDC EV and CMP-EV therapy

Given the augmented effect of CMP-EVs on cardiac function, we focused on myocardial immunohistochemical analysis to identify any differential morphometric changes in the heart between EV groups that may contribute to enhanced therapeutic efficacy by CMP-EVs. To assess the effect of EV therapy on remote zone interstitial fibrosis, we analyzed the remote regions of Masson’s trichrome stained LV sections at 28 days post-MI. Both CDC EVs and CMP-EVs demonstrated a significant and comparable reduction in interstitial fibrosis when compared with PBS control (PBS: 8.5 ± 0.4 vs. CDC-EV: 3.7 ± 0.8 vs. CMP-EV: 4.6 ± 0.8, % Interstitial Fibrosis) (n = 5 mice per group, mean ± SEM, *p<0.01 using one-way ANOVA with Tukey’s post-hoc test) (**Fig. 5**).

**Figure 5.**
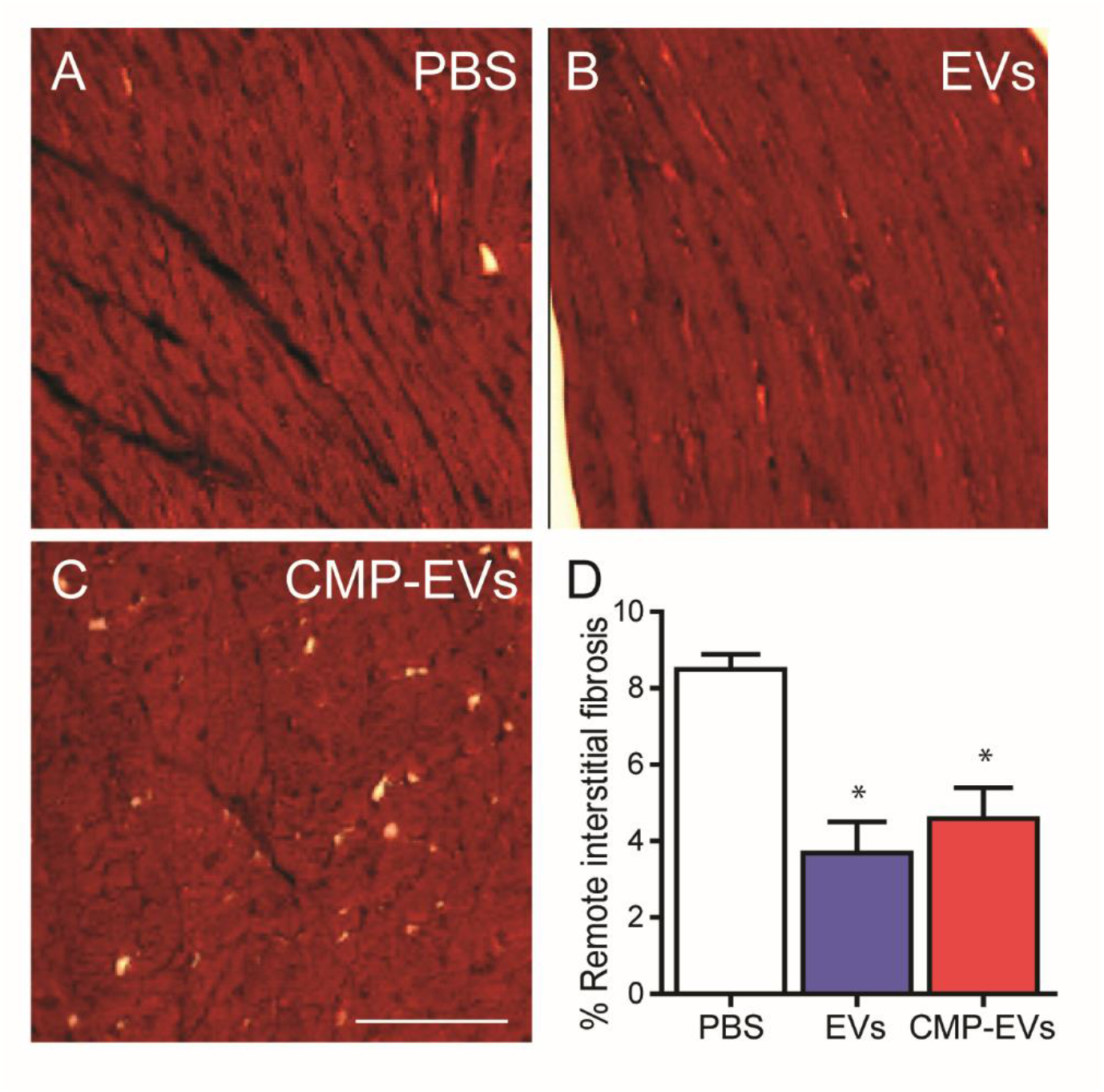
EV therapy modifies remote zone interstitial fibrosis post-MI. **(A-C)** Trichrome stained LV sections from the remote region 28 days post MI. **(B)** EVs and **(C)** CMP-EVs demonstrated a comparable reduction in interstitial fibrosis when compared with **(A)** PBS-vehicle control. n = 3 mice per group. Scale bar = 100 µm. *p<0.005 using a one-way ANOVA with Tukey’s post-hoc test. All data represented as mean ± SEM.

### CDC EVs and CMP-EVs reduce CD68^+^ macrophage infiltration post-MI

To investigate a potential differential effect of CDC EVs and CMP-EVs on modulating macrophage infiltration following ischemic injury, we analyzed CD68^+^ macrophage density in the infarct and remote zones 28 days post-MI. While macrophage density was highest in the infarct region and lowest in the remote zone for all treatment groups at 1 month, CDC EVs and CMP-EVs demonstrated a significant and comparable reduction in CD68^+^ macrophage density in the infarcted (PBS: 14 ± 2 vs. CDC-EV: 8 ± 1 vs. CMP-EV: 8 ± 2,% CD68 area) and remote myocardium (PBS: 4 ± 1 vs. CDC-EV: 1 ± 0.3 vs. CMP-EV: 1 ± 0.2,% CD68 area) when compared with PBS control (n=3 mice per group, mean ± SEM, *p<0.01 using one-way ANOVA with Tukey’s post-hoc test) (**Fig. 6**).

**Figure 6.**
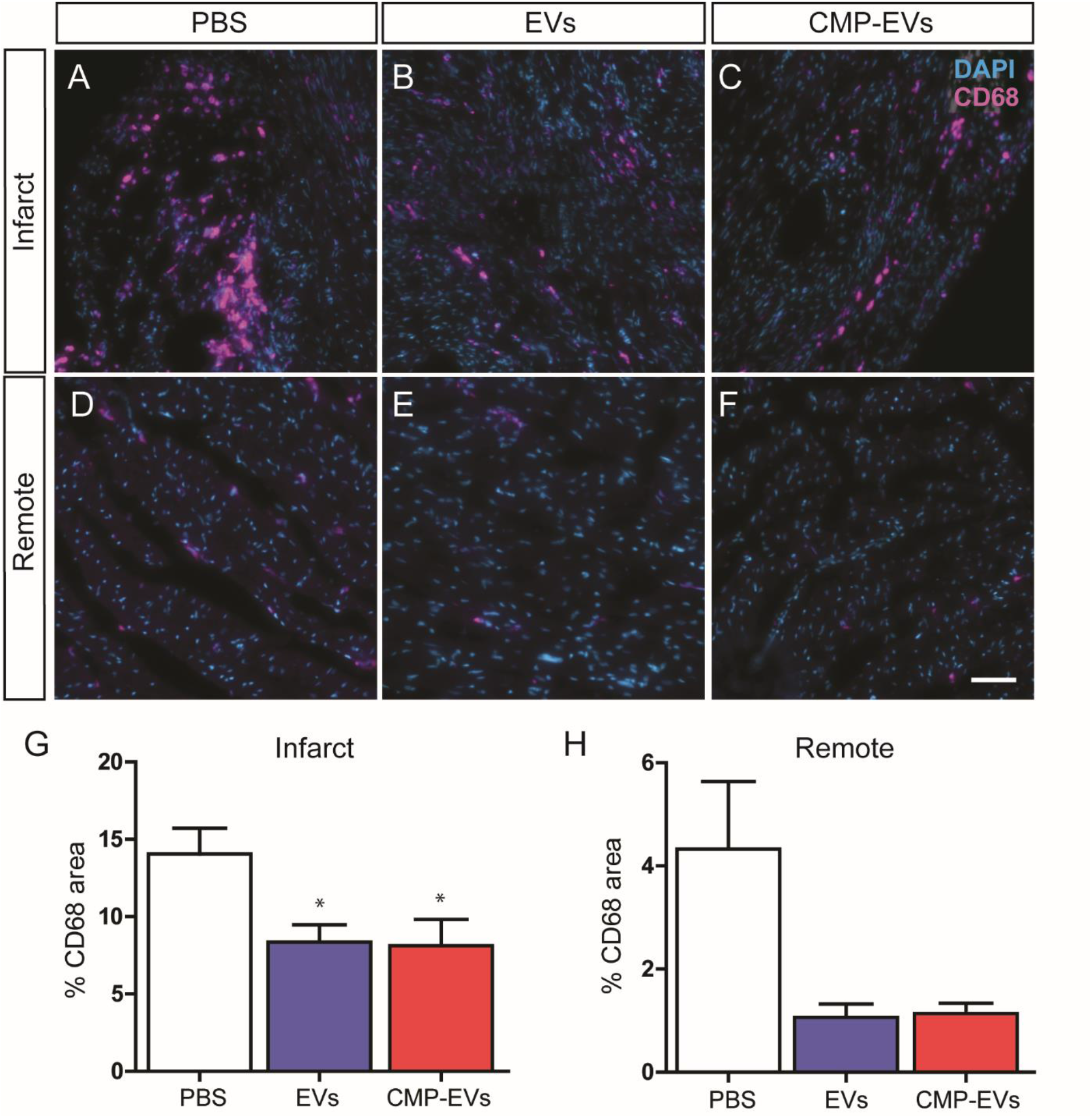
EV therapy reduces CD68^+^ macrophage density post-MI. CD68-stained sections from the **(A-C)** infarct and **(D-F)** remote zone 28 days post-MI. EVs and CMP-EVs demonstrated a comparable reduction in CD68+ macrophage density in the **(G)** infarcted and **(H)** remote myocardium. n = 3 mice per group. Scale bar = 50 µm. *p<0.05 using a one-way ANOVA followed by Tukey’s post hoc test. All data are represented as mean ± SEM.

### Targeted CMP-EVs demonstrate enhanced reduction of remote zone myocardial apoptosis

To examine the effect of EV therapy on apoptosis in the remote and infarct zone post-MI, we analyzed cleaved caspase-3-stained LV sections. The percentage of apoptotic cells was highest in the infarct region and lowest in the remote zone for all treatment groups at one month. While CDC EVs and CMP-EVs demonstrated a significant but equivalent reduction in the percentage of apoptotic cells in the infarct zone compared with PBS control (CDC-EV: 7 ± 1, CMP-EV: 7 ± 1, PBS: 11 ± 1, % Cleaved caspase-3+ cells), administration of CMP EVs further reduced the level of apoptosis beyond that of unmodified CDC EVs in the remote zone (CMP-EVs: 0.7 ± 0.1, CDC-EVs: 1.7 ± 0.2, PBS: 2.4 ± 0.1, % Cleaved caspase-3+ cells) (n=3 mice per group, mean ± SEM, *^,#^p<0.05 using one-way ANOVA with Tukey’s post-hoc test) (**Fig. 7**).

**Figure 7.**
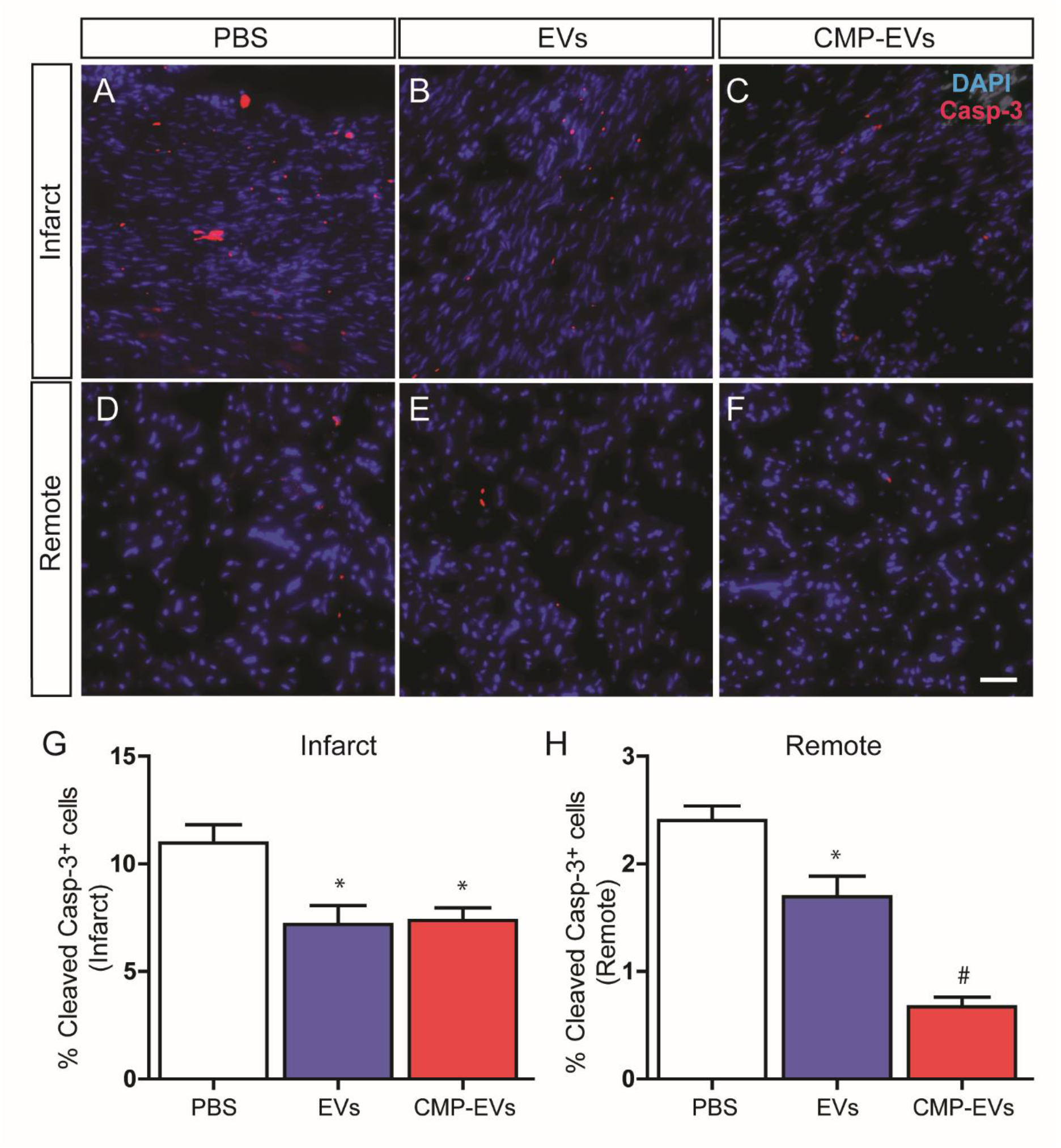
CMP-EVs augment EV salvage of remote zone apoptosis. Cleaved caspase-3-stained sections from the **(A-C)** infarct and **(D-F)** remote zone 28 days post-MI. **(G)** While EVs and CMP-EVs demonstrated a comparable reduction in apoptotic cells in the infarct zone, **(H)** administration of CMP-EVs further reduced the level of apoptosis beyond that of EVs in the remote zone. n = 3 mice per group. Scale bar = 50 µm. *p<0.05 (treatment vs. PBS control), ^#^p<0.05 (CDC-EVs vs. unmodified EVs) using a one-way ANOVA followed by Tukey’s post hoc test. All data are represented as mean ± SEM.

### EV treatment has a differential effect on rescuing cardiomyocyte vs. non-cardiomyocyte apoptosis in the remote zone

Based on the cell type-specific targeting of CMP-EVs to cardiomyocytes, we hypothesized that augmented improvement in remote zone apoptosis at one month following CMP-EV vs. EV treatment was secondary to rescue of myocyte cell death. To investigate this further, we quantified the number of cleaved caspase-3^+^/F-actin^+/-^ cells in the remote zone to identify the fraction of apoptotic cardiomyocytes (CMs) vs “non-cardiomyocytes” / total apoptotic cells in each treatment group (**Fig. 8a**). While there was no significant difference between PBS and EV treatment groups in the levels of “non-cardiomyocyte apoptosis” (which was the predominant source of cell death in all groups) (**Fig. 8b**), CMP-EV treatment resulted in a significant decrease in the percentage of apoptotic cardiomyocytes when compared with both PBS and unmodified EVs (CMP-EVs: 13% ± 2%, unmodified EVs: 19% ± 1%, PBS: 24% ± 8%) (n=3 mice per group, mean ± SEM, p<0.05 using one-tailed t-test) (**Fig. 8c**). When we quantified this as Casp-3^+^ CM/10^6^ CMs, we found that treatment with CMP-EVs significantly reduced the total number of apoptotic myocytes (4,006 ± 577/1×10^6^) compared with PBS and CDC EV treated hearts (PBS: 14,000 ± 2,290/1×10^6^, CDC EVs: 6,006 ± 577/10×10^6^) (n=3 mice per group, mean ± SEM, p<0.05 using one-way ANOVA with Tukey’s post-hoc test).

**Figure 8.**
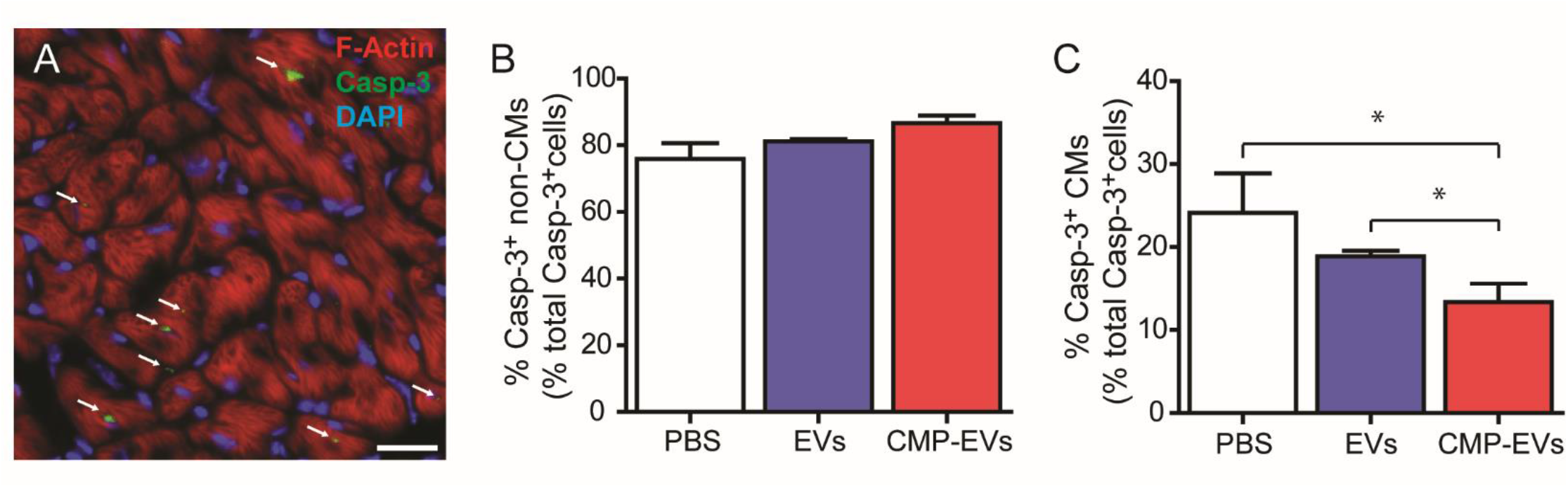
Cell-type specificity mediates CMP-EV reduction of remote zone apoptosis. **(A)** Cleaved caspase-3^+^ F-actin^+/-^ cells in the remote zone were quantified to identify the fraction of apoptotic cardiomyocytes vs. “non-cardiomyocytes”. **(B)** While there was no difference between PBS and EV treatment groups in the levels of non-cardiomyocyte apoptosis, **(C)** CDC-EV treatment resulted in a significant decrease in the percentage of apoptotic cardiomyocytes among total apoptotic cells when compared with both PBS and unmodified CDC EVs. n = 3 mice per group. *p<0.05 one-tailed t test of CMP-EVs vs. PBS and CMP-EVs vs. unmodified EVs. All data are represented as mean ± SEM.

## Discussion

The therapeutic efficacy of extracellular vesicles is largely defined by their biodistribution ^6^. In the current study, we demonstrate that engineering CDC EVs to target cardiac myocytes enhances their ability to augment systolic function and reduce infarct size following AMI. Similar to unmodified CDC EVs, surface engineered CMP-EVs decreased remote interstitial fibrosis and macrophage density at 30 days post-MI. Likely accounting for their augmented therapeutic efficacy, however, CMP-EVs demonstrated enhanced reduction of remote zone myocardial apoptosis which, when observed on a cellular level, appeared to be secondary to a specific effect on remote zone cardiomyocytes.

Several preclinical studies have highlighted the therapeutic potential of CDC EVs for the treatment of heart disease. CDC EVs have shown the capacity to reduce scar size ^8^, augment fibrosis ^12^, reduce cardiomyocyte apoptosis ^11^, promote angiogenesis ^7,9^, influence macrophage phenotype ^7,9^, and enhance cardiac function following MI and I/R ^13,14^. Many of these benefits can be attributed to specific miRNA cargo, such as miR-146a ^15^, miR-21 ^16^, and miR-181b ^8^, as well as Y RNA cargo ^5^ which has demonstrated a therapeutic effect post-MI. In addition to their efficacy in models of ischemic heart disease, CDC EVs have also shown promise in models of heart failure. For example, Bittle *et al.* demonstrated that treatment with CDC EVs resulted in attenuation of right ventricular dysfunction in the setting of acute pressure overload ^17^. Despite the success of CDC EVs in pre-clinical models, therapeutic efficacy appears to be dependent on myocardial retention of EVs, with prior studies demonstrating a need for intramyocardial delivery for infarct size reduction post-MI ^6^. We hypothesized that methods to improve cardiac retention of therapeutic EVs may further enhance treatment effect as well as reduce the dose necessary to elicit a therapeutic response, addressing a challenge in translating EV therapy to the clinic where scalability is of key importance.

Emerging evidence suggests that EVs can be engineered to hone their tropism by modification of their surface membrane, resulting in improved therapeutic efficacy ^18–21^. In the context of cardiovascular disease, a number of direct and indirect engineering strategies have been successfully employed in models of AMI ^22^ ^20^. Using bio-orthogonal chemistry, Zhu *et al.* directly expressed an ischemic myocardium-targeted peptide (IMTP) on the surface of hypoxia-conditioned MSC-EVs ^19^. Modified EVs demonstrated enhanced cardiac targeting post-MI which translated to improved function compared with control EVs ^19^. Chemical surface modifications and EV membrane cloaking have also been utilized to express an IMTP on CDC EVs resulting in enhanced efficacy in rat models of AMI. Similar to the indirect engineering strategy used in our study, Wang *et al.* genetically engineered MSCs to produce a Lamp2b-IMTP fusion protein which led to higher accumulation of EVs in the infarct zone and improved cardiac function post-MI ^23^. While both direct chemical and indirect genetic modification approaches generate modified EVs, indirect genetic engineering of parent cells may allow for the production of a more standardized and stable EV population ^24^. Employing an indirect surface engineering strategy to modify parent cells, we engineered CDCs to express a Lamp-2b-CMP fusion protein, resulting in the sustained production of cardiomyocyte targeting CMP-EVs. Importantly, these modifications did not affect the physical properties of CDC EVs (previously characterized in ^11^) as nanoparticle tracking analysis of CMP-targeted EVs showed a similar size distribution to unmodified EVs (mean diameter 123 nm) and cryo-TEM demonstrated an expected EV morphology of small, round vesicles with a clearly discernable lipid bilayer ^11^. microRNA content was comparable between groups as determined by similar levels of expression of characteristic CDC-EV miRs in CMP-EVs. Surface protein expression was also alike between unmodified CDC EVs and CMP-EVs, with both preparations positive for known EV markers CD63, CD81, ALIX, FLOT1, ICAM1, EpCam, ANXA5 and TSG101 and negative for any cellular contamination (by cis-Golgi marker GM130) ^11^. In addition, surface particle charge analysis demonstrated a similar negative zeta potential between groups ^11^. This in-depth characterization allowed us to attribute any differences in therapeutic efficacy and cellular responses to cardiomyocyte targeting and enhanced cardiac retention.

Cardiomyocytes represent an important therapeutic target in modulating the pathogenesis and progression of cardiac disease. Cardiac myocyte cell death is a sentinel event in the development of both ischemic and non-ischemic cardiomyopathies. Irreversible loss further contributes to additional myocyte dysregulation, generating a positive-feedback loop which potentiates the evolution of heart disease ^1^. To explore the efficacy of targeting therapeutic EVs to cardiomyocytes following cardiac injury, we engineered CDCs to generate EVs expressing a cardiomyocyte specific biding peptide on their surface. While our current study examines the therapeutic efficacy of CMP-EV treatment in a model of AMI, use of a cardiomyocyte specific peptide rather than an IMTP further extends their therapeutic applicability to non-ischemic heart disease. Additionally, the conserved nature of the peptide (between mice, swine and humans) reduces barriers to clinical translation.

Our study demonstrated several unexpected findings that may provide potential mechanistic insight into the cellular targets of CDC EVs in the cardiac milieu post-MI. Given the cardiomyocyte specific targeting of our CMP-EVs, we were surprised to see matched improvement between unmodified and targeted EVs in reducing remote interstitial fibrosis and macrophage density at 30 days post-MI. Cardiac fibrosis following AMI, while integral for the maintenance of structural integrity, has been associated with chronic deterioration of heart function ^25^. Prior studies have demonstrated an ability of CDC EVs to reprogram fibroblasts to generate EVs with similar therapeutic properties to CDC EVs ^12^. These results are consistent with other studies showing EV modification of their target cell secretome ^10^. As such, one mechanism by which CMP-EVs mediate remote zone interstitial fibrosis may be via indirect modification of the cardiomyocyte EV secretome to include anti-fibrotic miR cargo. In a similar vein, work by our group and others has highlighted a role of CDC-EVs in regulation of the immune response via direct effects on macrophage polarization ^7–9^. Our current study demonstrates that enhancing cardiomyocyte uptake of CDC-EVs does not diminish this effect. One hypothesis resulting from this work is that in addition to non-specific uptake by monocytes and macrophages, CMP-EVs may also modify the secretome of border and remote zone cardiomyocytes to indirectly modify macrophage properties post-MI. Additional research is needed to further elucidate these mechanisms and identify operant cargo, but it does not appear that either effect is directly responsible for the additional improvement in systolic function and infarct size conferred by CMP-EVs.

What is notably different between EV groups on a histological level is their effect on remote zone apoptosis 30 days post-MI. While CDC EVs and CMP-EVs demonstrated a significant but equivalent reduction in the percentage of apoptotic cells in the infarct zone compared with PBS, targeted CMP-EVs further reduced the level of remote zone myocardial apoptosis compared with unmodified CDC EVs. Additionally, when we quantified the number of cleaved caspase-3^+^/F-actin^+/-^ cells in the remote zone to identify the fraction of apoptotic cardiomyocytes vs non-cardiomyocytes as a percentage of total apoptotic cells, we found that CMP-EVs had the most significant effect on reducing cardiomyocyte apoptosis. This is likely secondary to the higher area and density of CMP-EV cardiac retention across the remote zone leading to enhanced delivery and uptake of cardioprotective miRs to remote zone cardiomyocytes. CDC EVs highly express a number of miRs which have been shown to have beneficial effects on cardiomyocyte viability. For example, miR-146a has been demonstrated to attenuate myocyte apoptosis and modulate autophagy through interactions with TATA-binding protein (TBP) associated factor 9b (TAF9b)/p53 as well as interleukin-1 receptor associated kinase1 (IRAK1) and tumor necrosis factor (TNF) receptor-associated factor 6 (TRAF6)/nuclear factor κΒ (NF-κΒ) ^26^. miR-21, also expressed highly by CDCs, has been shown to reduce myocardial apoptosis by modulating expression of programmed cell death 4 (PDCD4), FasL, and AKT pathways [47–49]. It is likely transfer of multiple miRs working in tandem that regulates cardiomyocyte viability.

Myocyte cell death occurs both acutely and chronically following myocardial infarction^27^. Therapeutics that result in small, sustained reductions in cell death, will likely have a cumulative effect in preservation of ventricular myocytes and prevention of heart failure. Our work highlights that targeting the remote myocardium, specifically remote cardiomyocytes, with engineered EVs holds promise as a viable therapeutic strategy for ischemic heart disease. Additional studies that couple EV cargo engineering with multi-cell type targeting and PK/PD assessment will further enhance the translational potential of EVs in ischemic and non-ischemic heart disease.

## Perspectives

### Competency in Medical Knowledge

While extracellular vesicles (EVs) have emerged as promising therapeutic vectors for cardiovascular disease, they are limited by their biodistribution. Engineering the surface of therapeutic EVs has recently been highlighted as a viable strategy to improve their pharmacokinetic properties and therapeutic efficacy, furthering their translational potential.

### Translational Outlook

Indirect surface engineering of CDC-EVs enhanced cardiomyocyte targeting and improved cardiac function in a preclinical model of acute myocardial infarction through rescue of remote zone apoptosis. This work highlights a role for targeting therapeutic EVs to cardiomyocytes following injury and serves as a proof-of-concept study for EV surface engineering in the treatment of CVD. Future studies are warranted to evaluate the pharmacokinetic and pharmacodynamic properties of engineered EVs in large animal models of heart disease prior to clinical translation.

## Acknowledgements

We would like to thank Dr. Joseph Spernyak and Roswell Park Comprehensive Cancer Center for the use of the IVIS Spectrum, supported by NIH grants P30CA016056 and S10OD016450.

## Abbreviations and Acronyms

AMI: acute myocardial infarction
CDC: cardiosphere-derived cell
CM: cardiomyocyte
CMP: cardiomyocyte specific peptide
EF: ejection fraction
EVs: extracellular vesicles
LV: left ventricle
MI: myocardial infarction
NTA: Nanoparticle Tracking Analysis

---

**Supplemental Figure 1:**
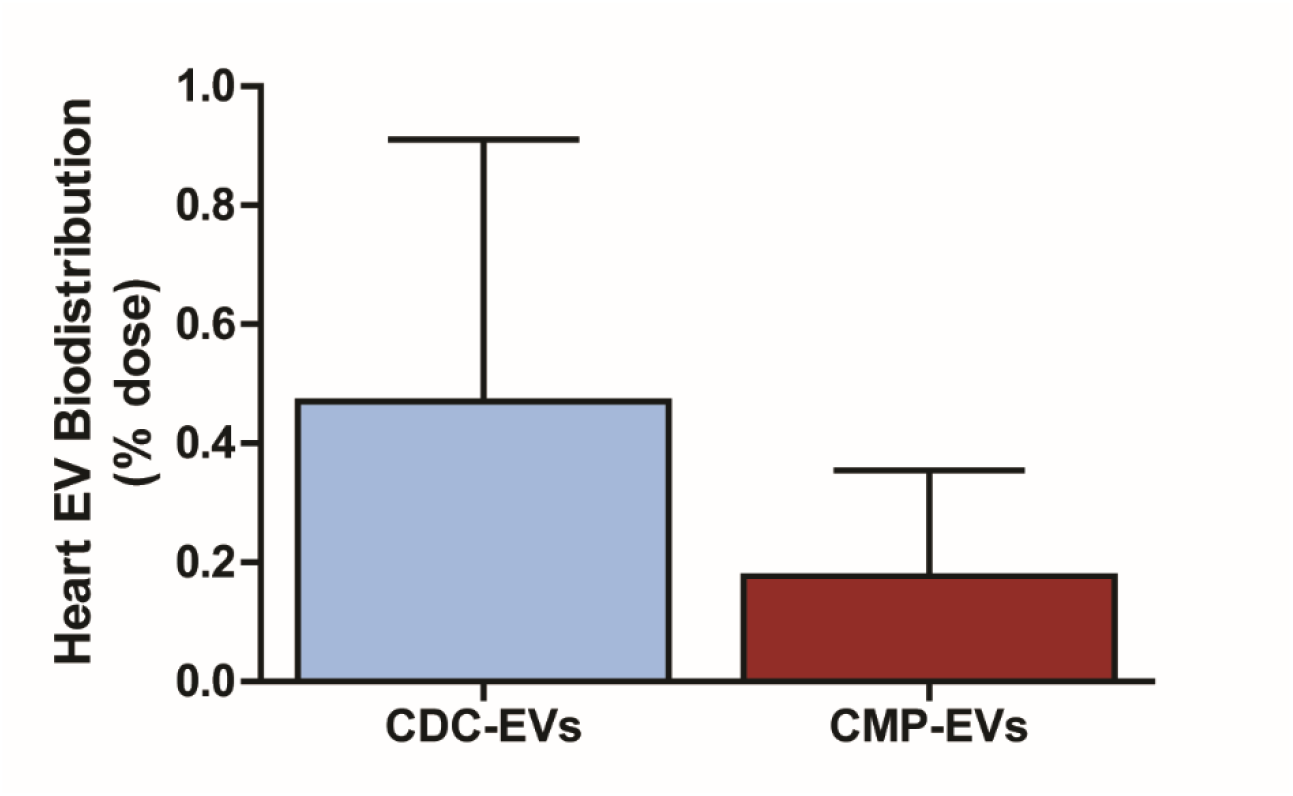
Cardiac biodistribution following systemic EV injection in healthy controls. DiR-labelled CDC EVs and CMP-EVs demonstrate no difference in cardiac targeting at 2 hrs following intravenous administration in healthy non-infarcted mice. n = 4-6 mice per group. p>0.05 using a two-tailed student’s t-test. All data are represented as mean ± SEM.

**Supplemental Figure 2:**
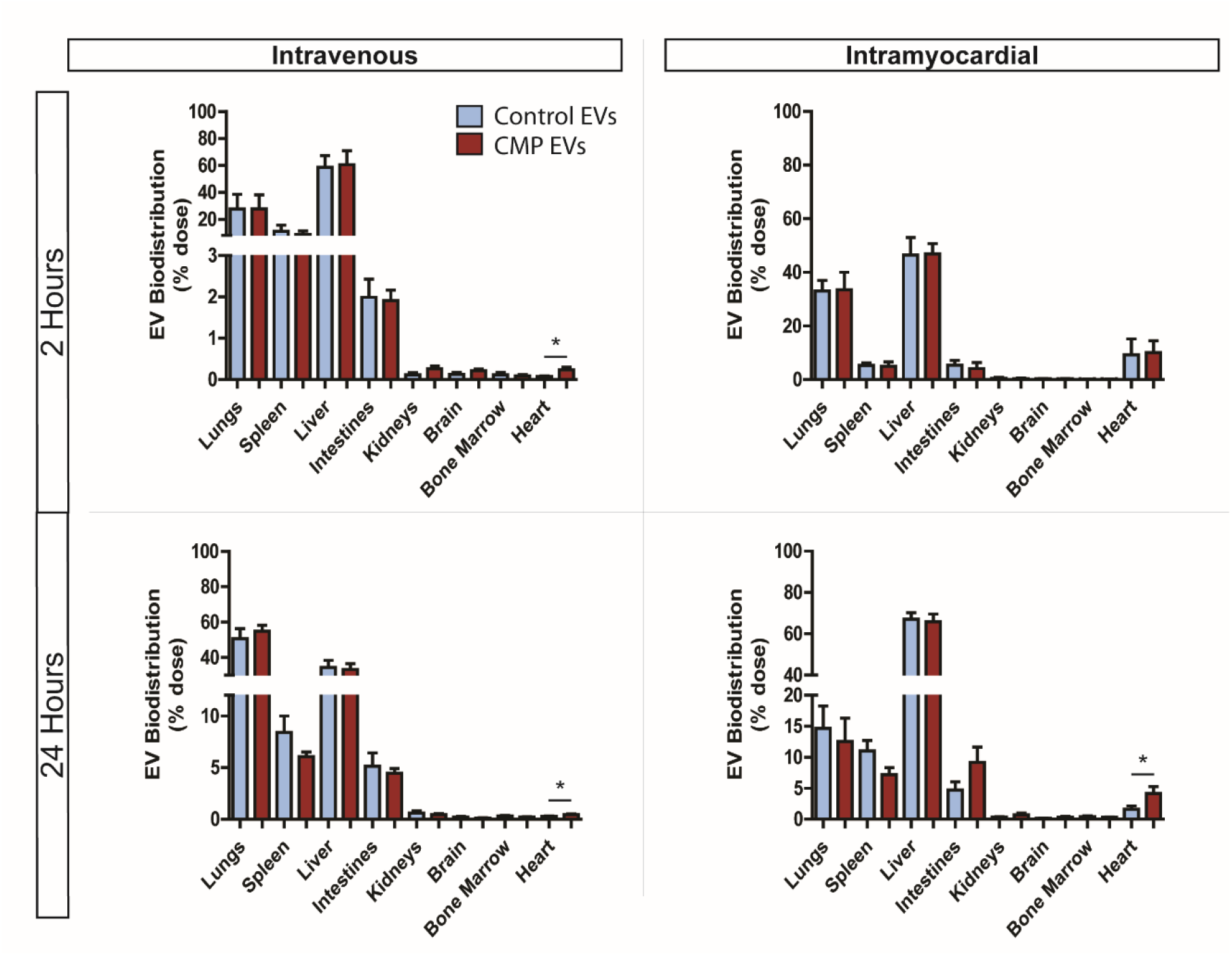
Comparison of total organ EV biodistribution and pharmacokinetics (PK) post-MI. Normalized DiD-stained EV biofluorescence signal as a percentage of total intravenous or intramyocardial injected dose at 2 and 24 hours. n = 5-9 mice per group. *p<0.05 using a two-tailed student’s t-test. All data are represented as mean ± SEM.

**Supplemental Figure 3:**
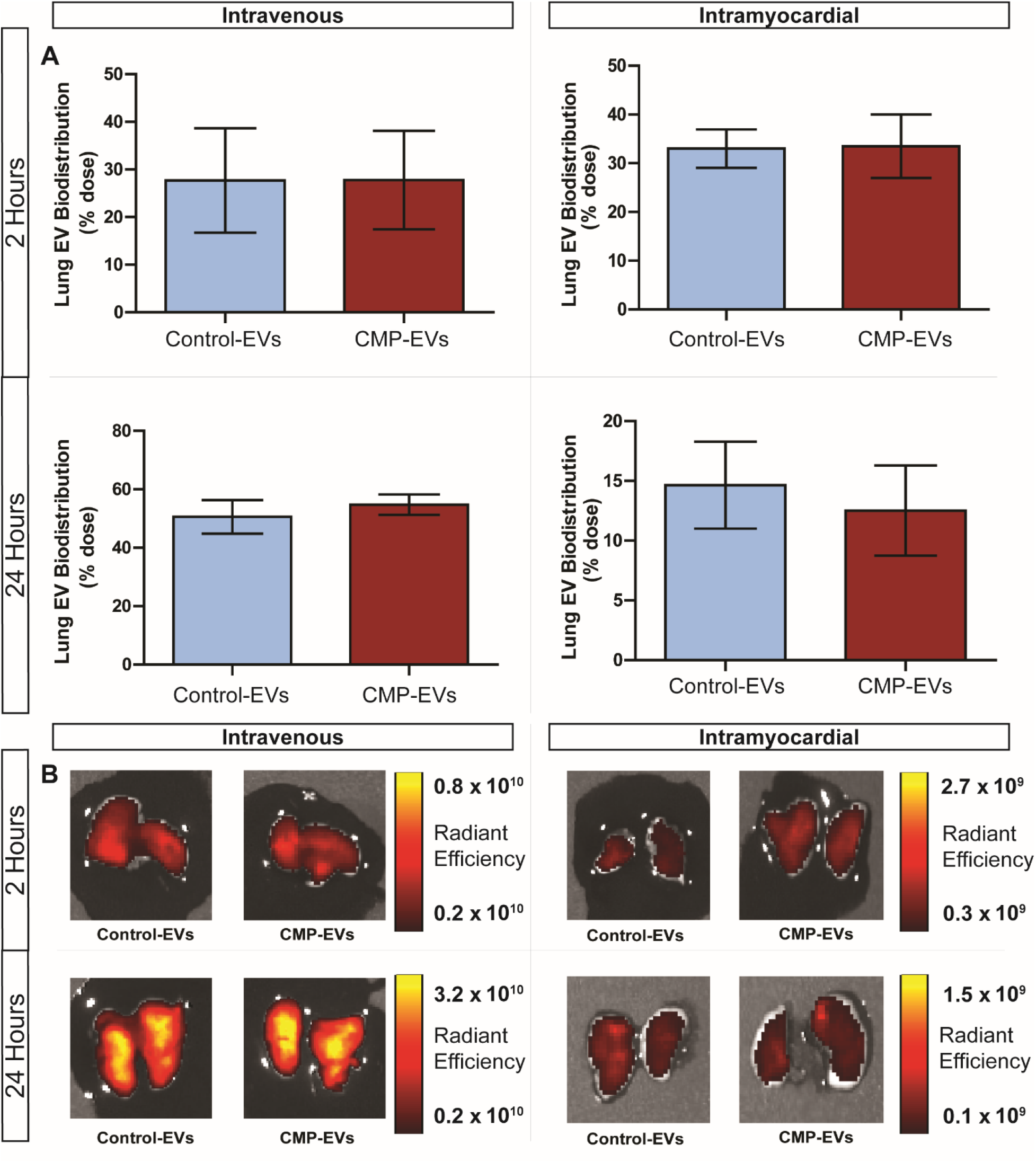
Lungs: EV biodistribution and PK post-MI. (**A**) Normalized DiD-stained EV biofluorescence signal in the lungs as a percentage of total injected dose at 2- and 24-hours following MI. (**B**) Representative IVIS images of the lungs demonstrate a similar retention of CDC EVs and CMP-EVs at each time point and administration route. n = 5-9 mice per group. p>0.05 using a two-tailed student’s t-test. All data are represented as mean ± SEM.

**Supplemental Figure 4:**
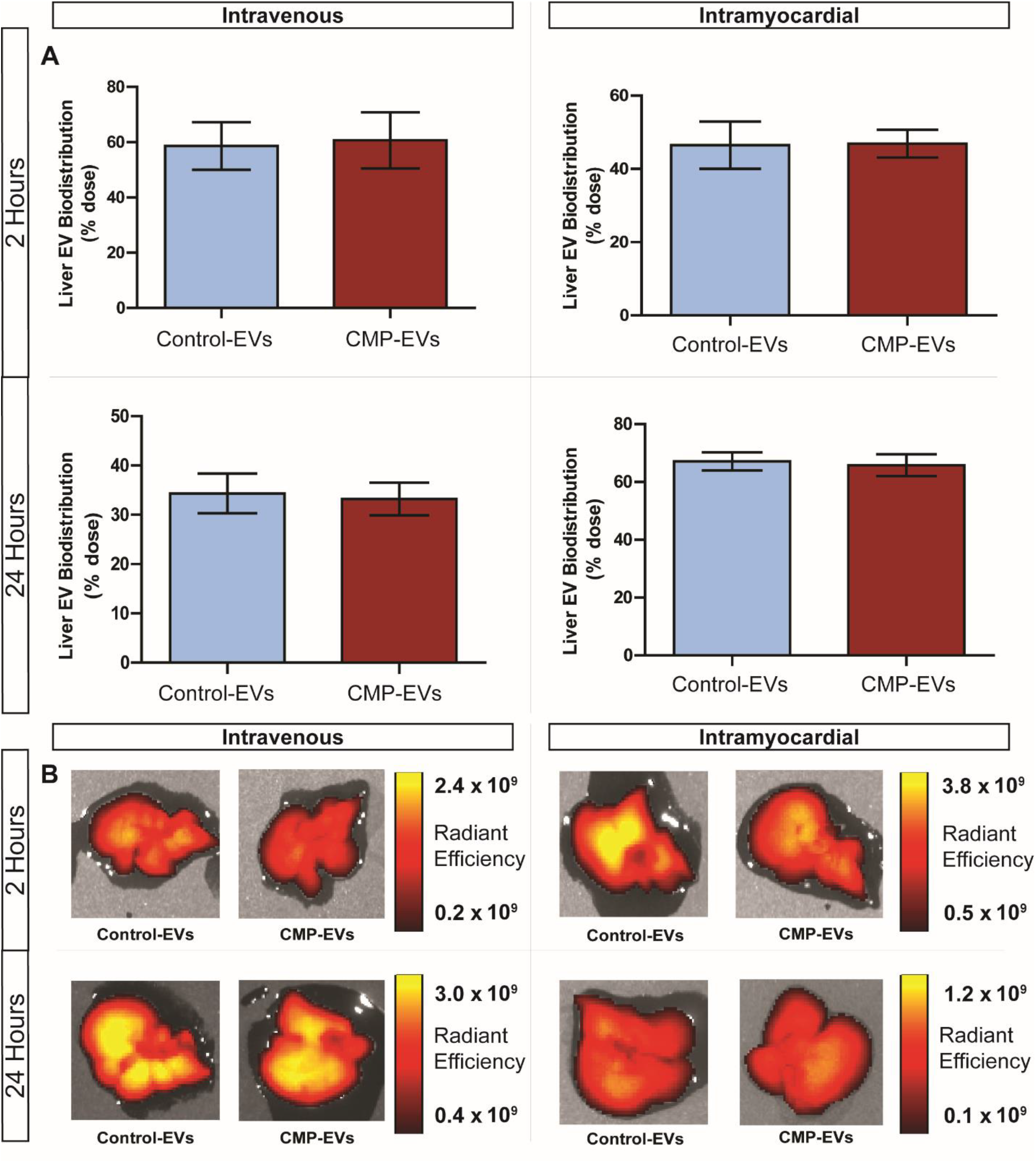
Liver: EV biodistribution and PK post-MI. (**A**) Normalized DiD-stained EV biofluorescence signal in the liver as a percentage of total injected dose at 2- and 24-hours following MI. (**B**) Representative IVIS images of the liver demonstrate a similar retention of CDC EVs and CMP-EVs at each time point and administration route. n = 5-9 mice per group. p>0.05 using a two-tailed student’s t-test. All data are represented as mean ± SEM.

**Supplemental Figure 5:**
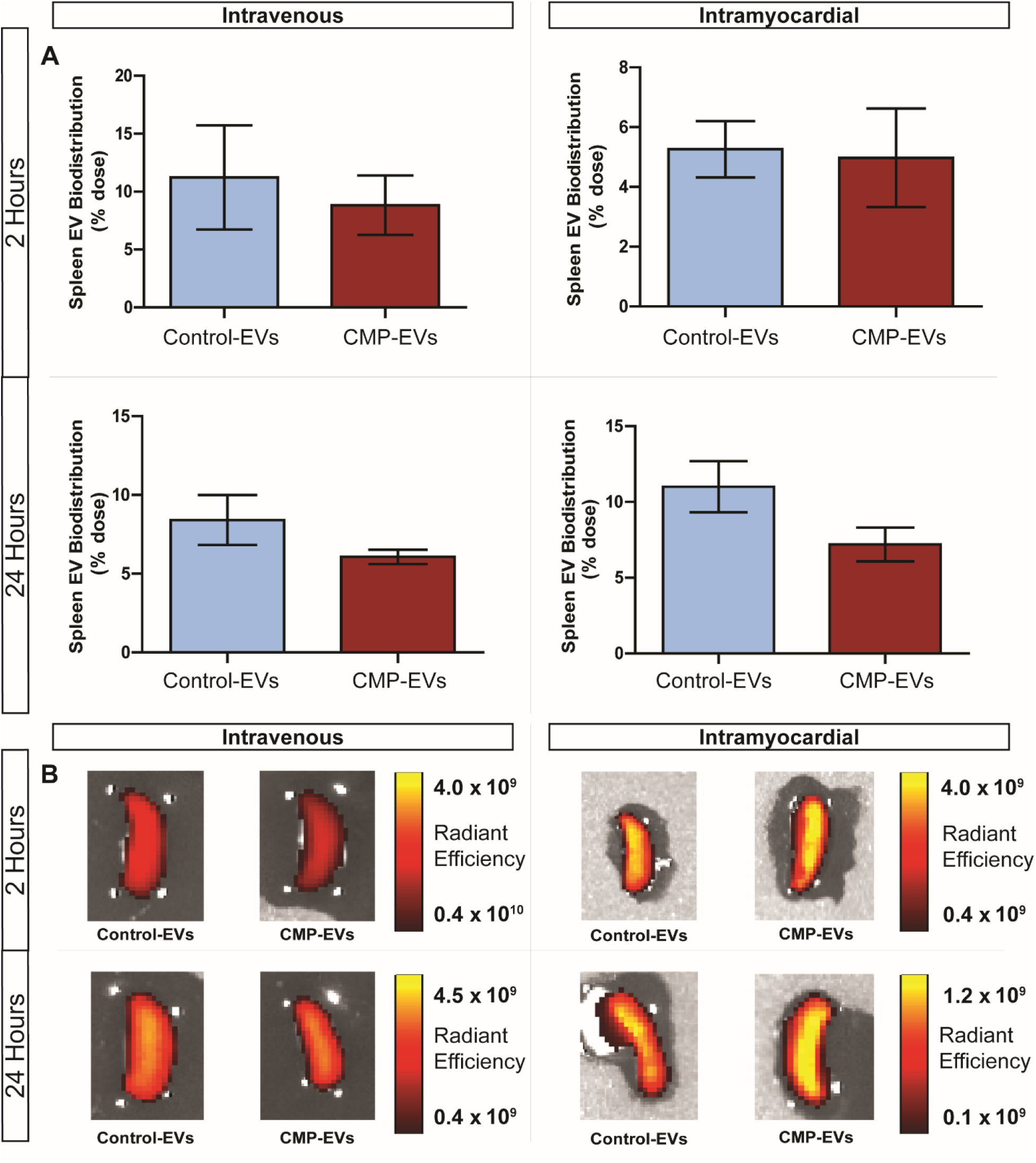
Spleen: EV biodistribution and PK post-MI. (**A**) Normalized DiD-stained EV biofluorescence signal in the spleen as a percentage of total injected dose at 2- and 24-hours following MI. (**B**) Representative IVIS images of the spleen demonstrate a similar retention of CDC EVs and CMP-EVs at each time point and administration route. n = 5-9 mice per group. p>0.05 using a two-tailed student’s t-test. All data are represented as mean ± SEM.

**Supplemental Figure 6:**
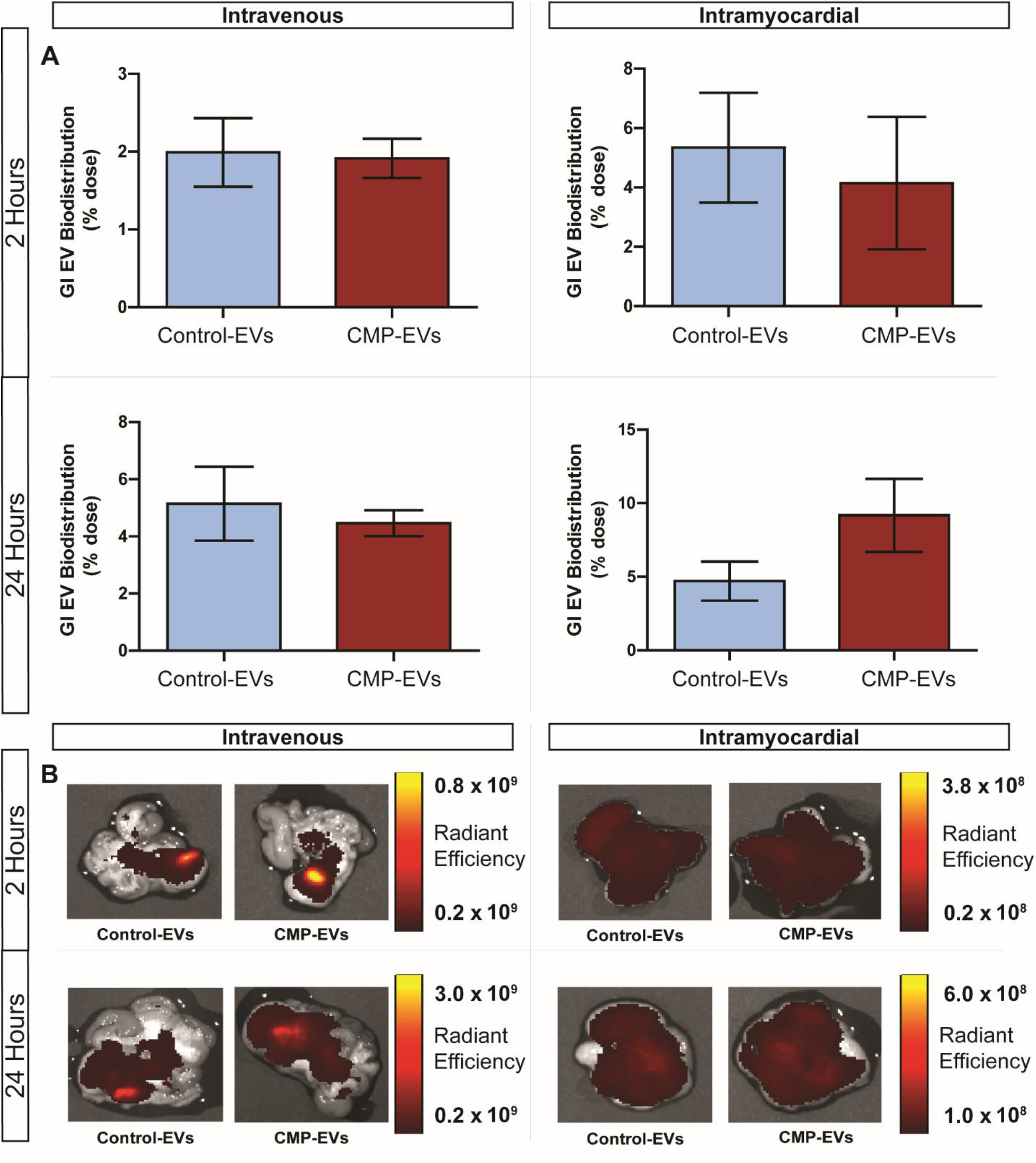
GI: EV biodistribution and PK post-MI. (**A**) Normalized DiD-stained EV biofluorescence signal in the GI track as a percentage of total injected dose at 2- and 24-hours following MI. (**B**) Representative IVIS images of the GI track demonstrate a similar retention of CDC EVs and CMP-EVs at each time point and administration route. n = 5-9 mice per group. p>0.05 using a two-tailed student’s t-test. All data are represented as mean ± SEM.

**Supplemental Figure 7:**
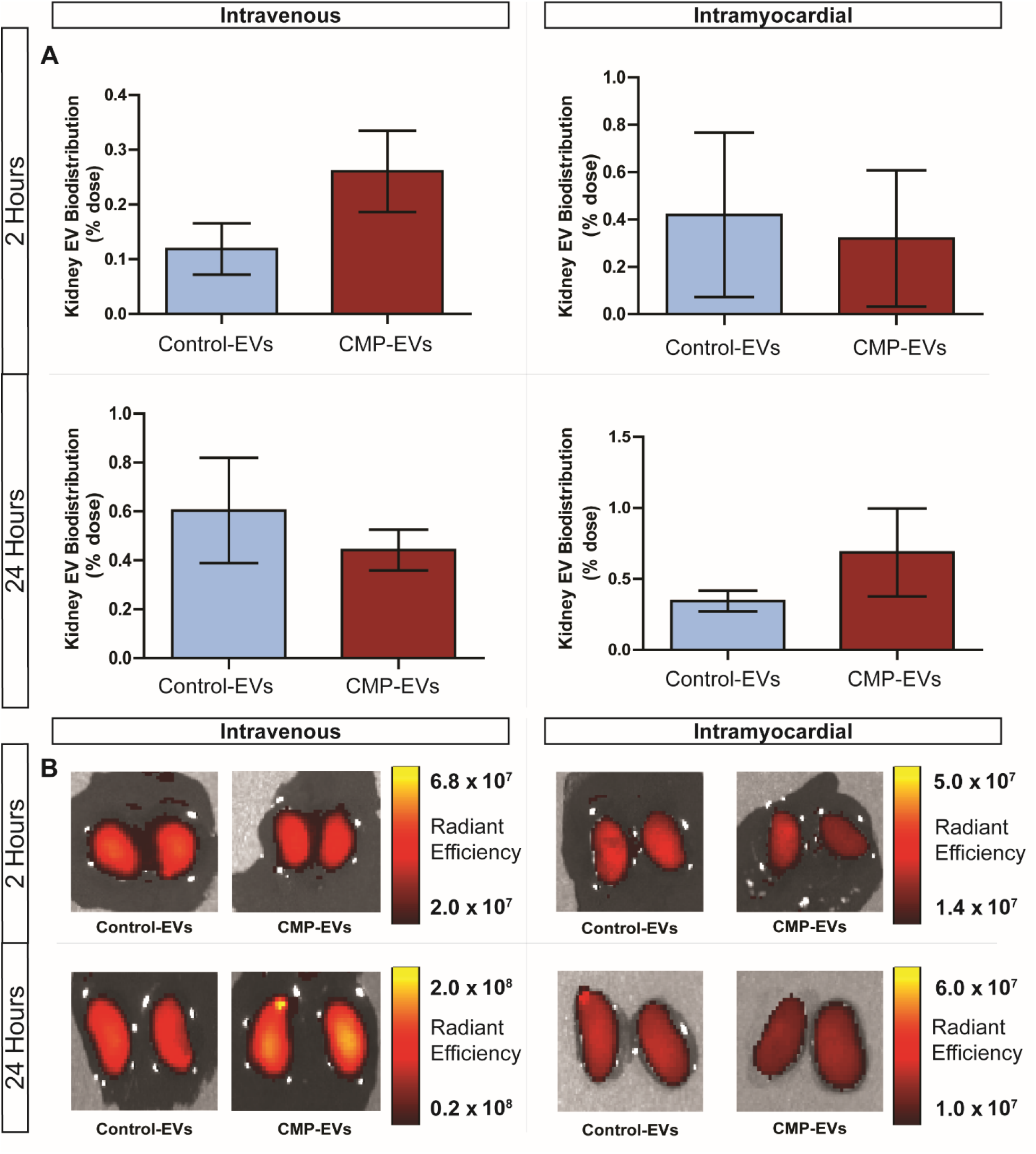
Kidneys: EV biodistribution and PK post-MI. (**A**) Normalized DiD-stained EV biofluorescence signal in the kidneys as a percentage of total injected dose at 2- and 24-hours following MI. (**B**) Representative IVIS images of the kidneys demonstrate a similar retention of CDC EVs and CMP-EVs at each time point and administration route. n = 5-9 mice per group. p>0.05 using a two-tailed student’s t-test. All data are represented as mean ± SEM.

**Supplemental Figure 8:**
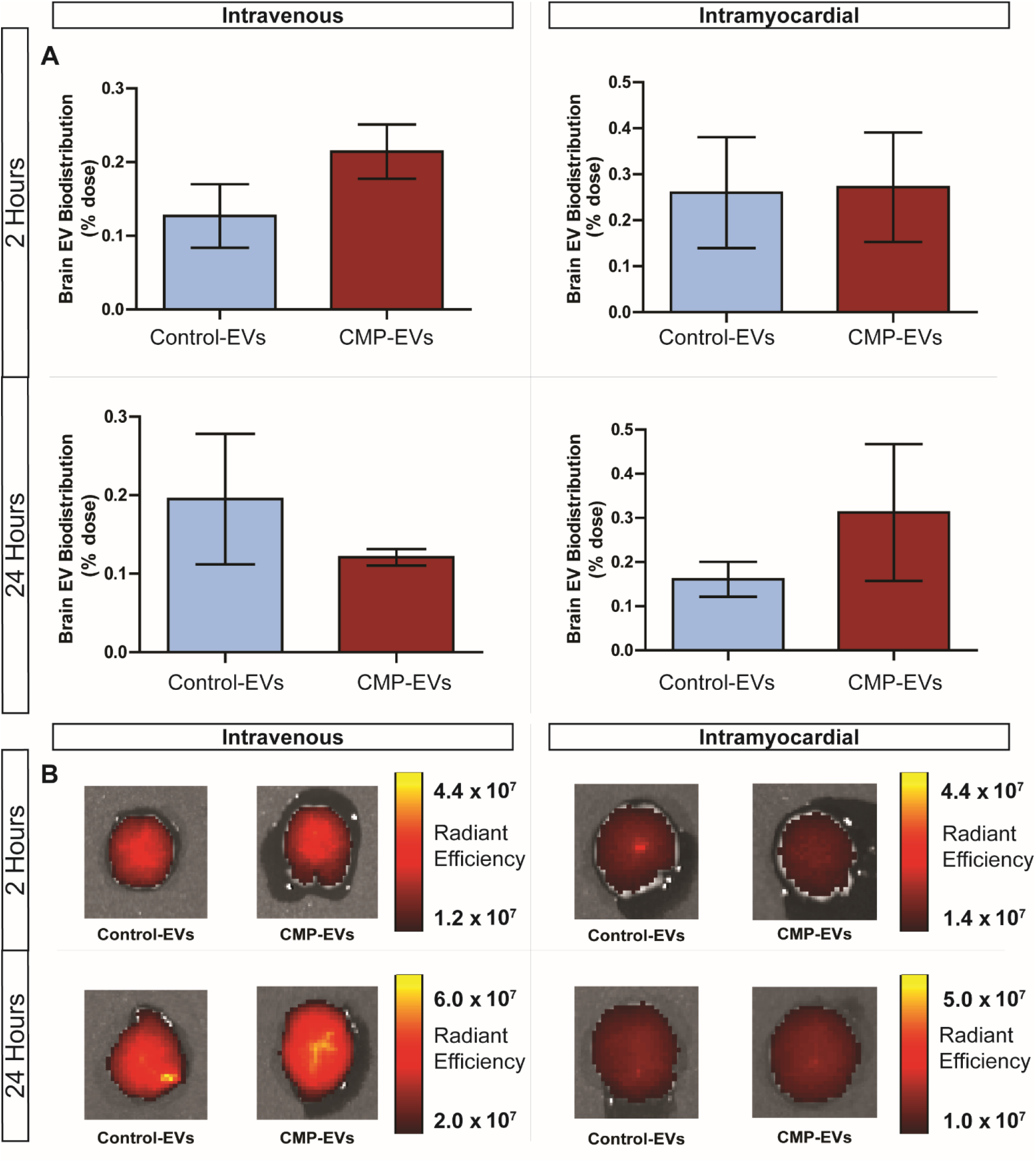
Brain: EV biodistribution and PK post-MI. (**A**) Normalized DiD-stained EV biofluorescence signal in the brain as a percentage of total injected dose at 2- and 24-hours following MI. (**B**) Representative IVIS images of the brain demonstrate a similar retention of CDC EVs and CMP-EVs at each time point and administration route. n = 5-9 mice per group. p>0.05 using a two-tailed student’s t-test. All data are represented as mean ± SEM.

**Supplemental Figure 9:**
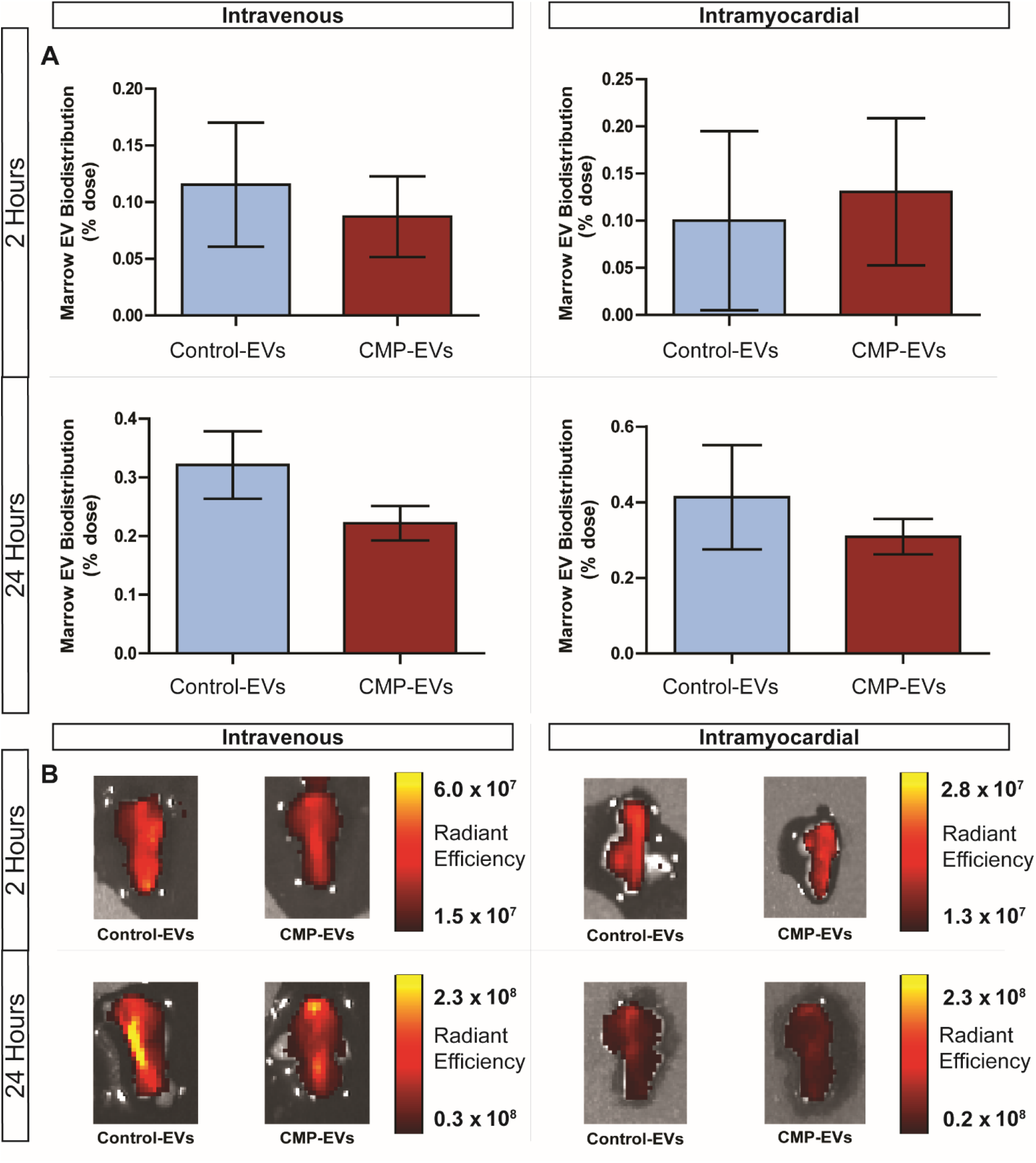
Bone Marrow: EV biodistribution and PK post-MI. (**A**) Normalized DiD-stained EV biofluorescence signal in the bone marrow as a percentage of total injected dose at 2- and 24-hours following MI. (**B**) Representative IVIS images of the bone marrow demonstrate a similar retention of CDC EVs and CMP-EVs at each time point and administration route. n = 5-9 mice per group. p>0.05 using a two-tailed student’s t-test. All data are represented as mean ± SEM.

**Supplemental Figure 10:**
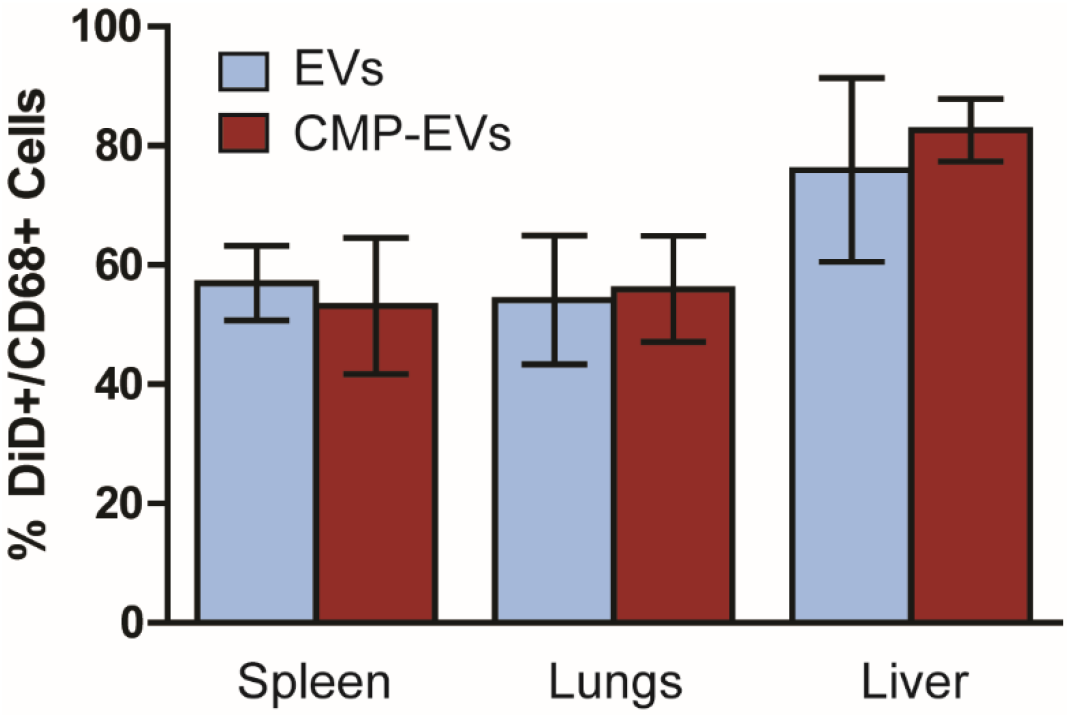
EVs are preferentially internalized by CD68+ macrophages of the liver, lungs and spleen. Cellular localization of DiD-labelled CDC EVs and CMP-EVs in the liver, lungs and spleen was assessed following EV IV delivery in health animals by flow cytometry. n = 3 mice per group. p>0.05 using a two-tailed student’s t-test on each organ. All data are represented as mean ± SEM.

